# DNA damage-induced senescence reshapes transcriptomic and functional landscape of human neural progenitor cells

**DOI:** 10.64898/2026.04.29.721535

**Authors:** Nisha V. Jose, Muhammad Z. K. Assir, Olivia Soper, Susan Janssens, Bettina Platt, Eunchai Kang

**Affiliations:** Institute of Medical Sciences, School of Medicine, Medical Sciences and Nutrition, University of Aberdeen, Aberdeen, AB25 2ZD, United Kingdom

**Keywords:** Aging, human neural progenitor cells, cellular senescence model, DNA damage response, hippocampal organoids, neurodegeneration

## Abstract

Ageing-related decline in hippocampal neurogenesis has been associated with cognitive impairment and neurodegenerative disease, yet experimentally tractable human models to study the underlying cellular and molecular mechanisms remain limited. Cellular senescence has emerged as a candidate driver of age-related tissue dysfunction, but its induction and consequences in human NPCs have not been well characterized. Here, we established a human in vitro model of NPC senescence using induced pluripotent stem cell-derived NPCs exposed to transient low-dose doxorubicin to activate the DNA damage response (DDR) while minimizing acute cytotoxicity. Doxorubicin-treated NPCs developed a stable senescent phenotype characterized by increased senescence-associated β-galactosidase activity, reduced proliferation, persistent DNA damage, and sustained induction of p21 and p16. Transcriptomic profiling revealed widespread senescence-associated remodeling, including activation of p53 and inflammatory programs and repression of cell cycle and DNA repair pathways. Senescent NPCs exhibited apoptosis resistance despite transcriptional priming of apoptotic pathways and underwent mitochondrial remodeling with a shift towards oxidative metabolism. In parallel, they acquired a senescence-associated secretory phenotype enriched in inflammatory, TGFβ-related and pro-angiogenic factors, and conditioned media from these cells promoted angiogenesis in vascular organoids. Importantly, key senescence-associated features were recapitulated in human hippocampal organoids, confirming the robustness of this paradigm in a three-dimensional neural context. Together, these findings establish a tractable human model of DDR-driven NPC senescence and identify senescence as a mechanism linking genotoxic stress to impaired progenitor function, metabolic rewiring, and paracrine niche remodeling relevant to hippocampal ageing and neurodegeneration.

## 1. Introduction

Ageing is the primary risk factor for neurodegenerative disorders, including Alzheimer’s disease and related dementias (De Strooper and Karran 2016, Hou et al. 2019). At the cellular level, ageing is characterized by the progressive accumulation of molecular damage, leading to functional decline and the emergence of key hallmarks such as genomic instability, altered metabolism, and cellular senescence (Lopez-Otin et al. 2013, Lopez-Otin et al. 2023). While senescence has been extensively studied in proliferative peripheral tissues (Campisi 2013, Gorgoulis et al. 2019, van Deursen 2014), its role within the human brain remains less well defined, particularly in neural progenitor cell (NPC) populations that retain proliferative capacity across the lifespan.

The adult hippocampus is one of the few regions of the mammalian brain that supports neurogenesis throughout life (Bond, Ming and Song 2015). NPCs residing in the dentate gyrus generate new neurons that contribute to learning, memory, and cognitive flexibility (Aimone et al. 2014, Deng, Aimone and Gage 2010, Toda and Gage 2018). However, hippocampal neurogenesis declines with age, and this reduction has been linked to cognitive impairment and neurodegenerative disease. Recent studies have provided evidence that adult hippocampal neurogenesis persists in humans but declines markedly with age, although the extent of this process remains debated (Boldrini et al. 2018, Moreno-Jimenez et al. 2019). More recent analyses, including single-cell transcriptomic approaches, support the presence of rare neural progenitor and immature neuronal populations in the adult human hippocampus, while highlighting substantial age-related reductions in their abundance (Dumitru et al. 2025, Franjic et al. 2022, Terreros-Roncal et al. 2021, Tobin et al. 2019). Importantly, analyses of human hippocampal tissue across ageing and neurodegenerative disease demonstrate that preservation of neurogenesis markers is associated with maintained cognitive function, suggesting that sustained neural progenitor activity may underlie successful brain ageing (Terreros-Roncal et al. 2021). Consistent with this, recent multi-omics profiling further shows that age-related cognitive decline is accompanied by disruption of neurogenic programs, whereas individuals with preserved cognition retain higher levels of neural progenitor activity, highlighting a potential link between NPC dysfunction and brain ageing (Disouky et al. 2026). The mechanisms underlying this decline remain incompletely understood, but intrinsic changes within NPCs are thought to play a central role.

NPCs maintain a tightly regulated balance between self-renewal and differentiation (Ming and Song 2011). During ageing, these cells exhibit reduced proliferative capacity, altered lineage commitment, and impaired regenerative potential (Encinas et al. 2011, Kempermann et al. 2018). Cellular senescence has emerged as a key mechanism that may contribute to these changes. Senescent cells undergo stable cell cycle arrest and acquire distinct phenotypic features, including resistance to apoptosis and the secretion of pro-inflammatory factors collectively termed the senescence-associated secretory phenotype (Campisi 2013, Gorgoulis et al. 2019). In the brain, accumulation of senescent cells may disrupt the neurogenic niche and impair tissue homeostasis; however, the extent and functional consequences of senescence in human NPCs remain poorly characterized.

A major driver of cellular senescence is the accumulation of DNA damage and the activation of the DNA damage response (DDR). Persistent DDR signaling can trigger a stable senescent state, linking genomic instability to cellular ageing (d’Adda di Fagagna 2008, Gorgoulis et al. 2019). In neural systems, increased DNA damage has been observed during ageing and in neurodegenerative conditions, suggesting that DDR-mediated senescence may contribute to impaired neurogenesis (Lu et al. 2004, Madabhushi, Pan and Tsai 2014). Nevertheless, mechanistic studies in human NPCs have been limited by the lack of well-controlled experimental models that can reliably induce senescence without confounding effects such as acute cytotoxicity.

To address this, we established a human in vitro model of NPC senescence using induced pluripotent stem cell (iPSC)-derived NPCs. Senescence was induced through controlled activation of the DDR using doxorubicin, a DNA-damaging agent widely employed to model genotoxic stress (Gewirtz 1999, te Poele et al. 2002). The treatment conditions were optimized to generate a stable senescent phenotype while minimizing cytotoxicity, enabling the investigation of senescence-specific effects in human NPCs.

Using this two-dimensional (2D) system, we validated key hallmarks of senescence and characterised the associated transcriptomic and functional changes. To extend the physiological relevance of our findings, we further validated senescence-associated features in a three-dimensional (3D) hippocampal organoid model, demonstrating that this phenotype can be recapitulated in a more complex, tissue-like context.

Collectively, this study establishes a robust and scalable human model of NPC senescence and provides a framework to investigate how DNA damage-driven senescence contributes to progenitor dysfunction and tissue-level changes relevant to brain ageing.

## 2. Results

### 2.1 Doxorubicin induces a senescence-like transcriptional program in neural progenitor cells

To induce senescence in neural progenitor cells (NPCs), cultures were treated with doxorubicin (Dox), a topoisomerase II–inhibiting chemotherapeutic that generates DNA double-strand breaks and blocks replication. Dox has been shown previously to trigger senescence across multiple cell types, including endothelial cells, cardiac fibroblasts, and cardiac progenitor populations (Gao et al. 2023, Zhang et al. 2021). NPCs were exposed to 25 nM Dox for 24 hours, followed by three-day drug-free period to allow the establishment of persistent senescence-associated phenotypes (**Fig. 1A**).

**Figure 1.**
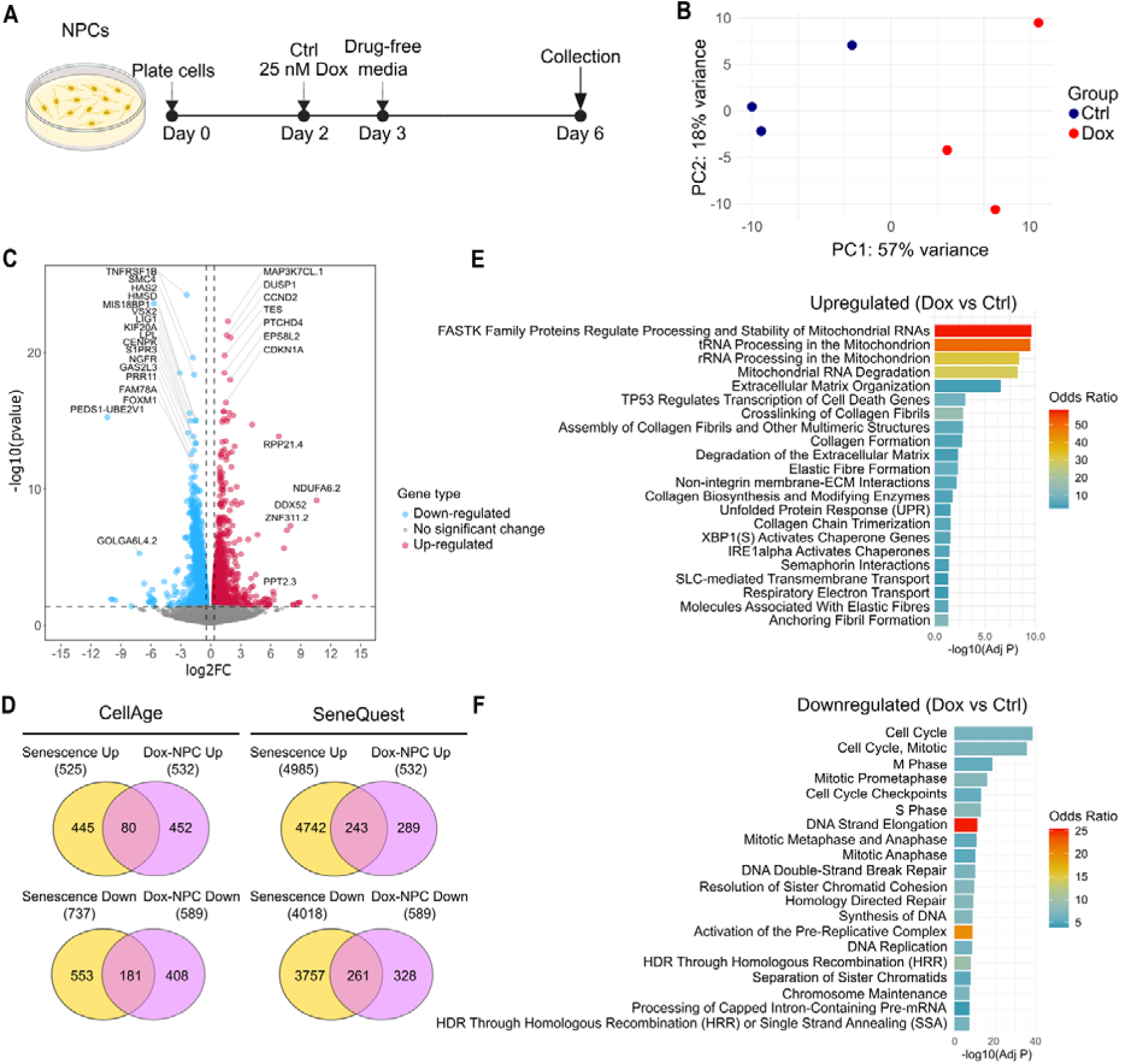
Doxorubicin induces a senescence like transcriptional program in NPCs. **(A)** Schematic of the experimental workflow. NPCs were treated with 25 nM doxorubicin (Dox) for 24 hours. Following drug removal on day 3, cells were maintained in drug-free medium for an additional four days to allow senescence-associated phenotypes to develop. Cells were harvested on day 6 for downstream analyses. **(B)** Principal component analysis (PCA) of bulk RNA-seq data shows clear separation between Ctrl and Dox-treated NPCs along PC1, which accounts for 57% of total variance. PC2 explains an additional 18% and captures within-group heterogeneity. Each point represents an individual sample. **(C)** Volcano plot displaying log_2_ fold change (x-axis) versus -log_10_ FDR-adjusted *p-*value (y-axis). Upregulated genes are shown in red, downregulated genes in blue, and nonsignificant features in grey. Dashed lines mark thresholds for statistical significance (FDR-adjusted *p* < 0.05) and effect size (log_2_FC ≥ 0.4). **(D)** Venn diagram illustrating overlap between differentially expressed genes in Dox-treated NPCs and senescence-associated genes in CellAge and SeneQuest databases. **(E–F)** Reactome pathway enrichment analysis for upregulated genes (**E**) and downregulated genes (**F**) in Dox-treated NPCs relative to controls. Pathways are ranked by adjusted *p-*value, and color intensity reflects Odds ratio calculated using Enrichr.

Bulk RNA sequencing revealed that Dox-treated NPCs adopt a transcriptional state markedly distinct from untreated controls. Principal component analysis (PCA) demonstrated clear segregation of samples along PC1, which accounted for 57% of total variance, while PC2 (18% variance) captured within-condition heterogeneity (**Fig. 1B**). Differential expression analysis identified substantial transcriptomic remodeling following Dox exposure, with 532 significantly upregulated and 589 significantly downregulated genes relative to controls (**Fig. 1C, Table S1**).

*CDKN1A* (p21) was among the significantly upregulated genes, consistent with its role as a canonical effector of p53-mediated cell cycle arrest and a well-established marker of senescence. Additional upregulated genes, including *CCND2*, *DUSP1*, *PTCHD4*, as well as numerous genes linked to mitochondrial function and RNA metabolism, indicate activation of stress response pathways and metabolic reprogramming characteristic of senescent cells. In contrast, downregulated genes were highly enriched for mitotic, and proliferation associated regulators, including *FOXM1*, *CENPF*, *KIF20A*, *MIS18BP1*, *GAS2L3*, and *PRR11*. The coordinated repression of these cell cycle drivers, alongside robust induction of p21, supports a transition from a proliferative NPC state to stable cell cycle arrest and a senescent phenotype (**Fig. 1C**, **Table S1**).

To evaluate whether Dox-treated NPCs activate established senescence gene programs, we compared the differentially expressed genes with curated senescence signatures from two complementary resources: CellAge, a manually curated database of genes experimentally shown to promote or inhibit cellular senescence (Avelar et al. 2020), and SeneQuest, an integrative knowledge base compiling senescence-associated genes and regulatory pathways from the literature (Gorgoulis et al. 2019). Within CellAge dataset, 80 of 532 upregulated genes and 181 of 589 downregulated genes in Dox-treated NPCs overlapped with genes reported to be up- or downregulated in cellular senescence, respectively (**Fig. 1D**). A similar pattern was observed using SeneQuest, with 243 of 532 upregulated and 261 of 589 downregulated genes matching known senescence-associated transcriptional changes (**Fig. 1D**, **Table S2**). Importantly, these overlaps showed high concordance in direction, with 97.8% of shared genes in CellAge and 72.5% in SeneQuest exhibiting consistent regulation with reported senescence signature (**Fig. S1A**). Together, these findings demonstrate that Dox-treated NPCs undergo transcriptional rewiring that closely mirrors established senescence programs

To further characterize the pathways altered during Dox-induced senescence, we performed Reactome pathway enrichment analysis on genes differentially expressed between Dox-treated and control NPCs. Upregulated genes were strongly enriched for mitochondrial RNA metabolic processes, including mitochondrial RNA processing and stabilization, maturation of mitochondrial rRNAs and tRNAs, and mitochondrial RNA turnover (**Fig. 1E**). These changes are consistent with the extensive mitochondrial remodeling observed in mitochondrial dysfunction–associated senescence (MiDAS) (Wiley et al. 2016). Dox-treated NPCs also showed robust activation of p53-dependent transcriptional programs, encompassing pathways involved in DNA damage responses and regulated cell death (**Fig. 1E**). In parallel, we observed enrichment of endoplasmic reticulum (ER) stress and unfolded protein response (UPR) pathways, including IRE1α-mediated chaperone activation and XBP1s-driven protein folding programs (**Fig. 1E**). These pathways reflect proteostasis disruption, a well-established contributor to senescence induction and maintenance. Additionally, we detected broad activation of extracellular matrix (ECM) remodeling pathways, including collagen biosynthesis and crosslinking, elastic fiber assembly, ECM degradation, non-integrin ECM interactions, and semaphorin signaling (**Fig. 1E**). This ECM remodeling is a hallmark feature of the senescence-associated secretory phenotype (SASP), which reshapes the surrounding microenvironment.

Conversely, genes downregulated in Dox-treated NPCs were strongly enriched for pathways governing cell cycle progression and DNA replication (**Fig. 1F**). Key suppressed modules included S-phase and M-phase regulation, cell cycle checkpoints, DNA strand elongation, pre-replicative complex activation, and core DNA replication machinery, indicating a loss of proliferative capacity. High-fidelity DNA repair pathways, such as homology directed repair, double strand break repair, and sister chromatid cohesion maintenance, were also markedly depleted (**Fig. 1F**), consistent with sustained DNA damage and replication stress, both characteristic hallmarks of senescence.

We also observed a substantial subset of NPC-specific changes compared to the CellAge and SeneQuest datasets (**Fig. S1B, C**). To further characterise these NPC-specific changes, we performed Reactome pathway enrichment analysis on 260 differentially upregulated genes that did not overlap with upregulated genes in the CellAge or SeneQuest datasets. These upregulated genes were strongly enriched for mitochondrial RNA processing and degradation pathways. While this enrichment is consistent with features commonly observed in senescence, a subset of these genes may reflect NPC-specific changes (**Fig. S1D**). Intriguingly, Reactome pathway and WikiPathways enrichment analyses of 263 NPC-specific downregulated genes revealed significant suppression of cholesterol metabolism–associated pathways including cholesterol biosynthesis and cholesterol synthesis disorders (**Fig. S1E, F**). These NPC-specific changes highlight the importance of studying senescence in a cell type–specific context.

Together, these pathway-level alterations in transcriptomic analyses indicate that Dox exposure drives NPCs into a robust senescence-like state characterized by coordinated mitochondrial and ER stress, activation of p53 signaling, repression of cell cycle and DNA repair programs, SASP-associated ECM remodeling, and the emergence of distinct NPC-specific transcriptional signatures.

### 2.2 Doxorubicin induces senescence like-morphological remodeling and coordinated cell-cycle suppression in NPCs

To confirm the maintenance of NPC identity upon Dox treatment, we first examined canonical NPC markers. Immunocytochemistry revealed robust expressions of SOX2, PAX6, and Nestin (**Fig. 2A**), confirming preservation of an undifferentiated NPC state throughout the experimental timeline. Dox exposure induced pronounced morphological changes characteristic of stress-associated senescence. Brightfield imaging of Dox-treated NPCs demonstrated marked cellular enlargement with increased flattening and spreading compared to the compact morphology of untreated controls (**Fig. 2B**). These observations were further supported by immunofluorescence staining with Nestin and DAPI, followed by confocal imaging and 3D reconstruction to resolve cellular and nuclear architecture (**Fig. 2C, Fig. S1G**). Quantitative analysis revealed significant increases in nuclear and cytoplasmic areas and perimeters in Dox-treated NPCs, consistent with nuclear enlargement and overall cell expansion (**Fig. 2D-G**). Collectively, these features, including enlarged nuclei, expanded cytoplasmic volume, and altered cell shape, are consistent with well-established senescence phenotypes across diverse cell types and indicate that NPCs mount a senescence-like response following Dox treatment.

**Figure 2.**
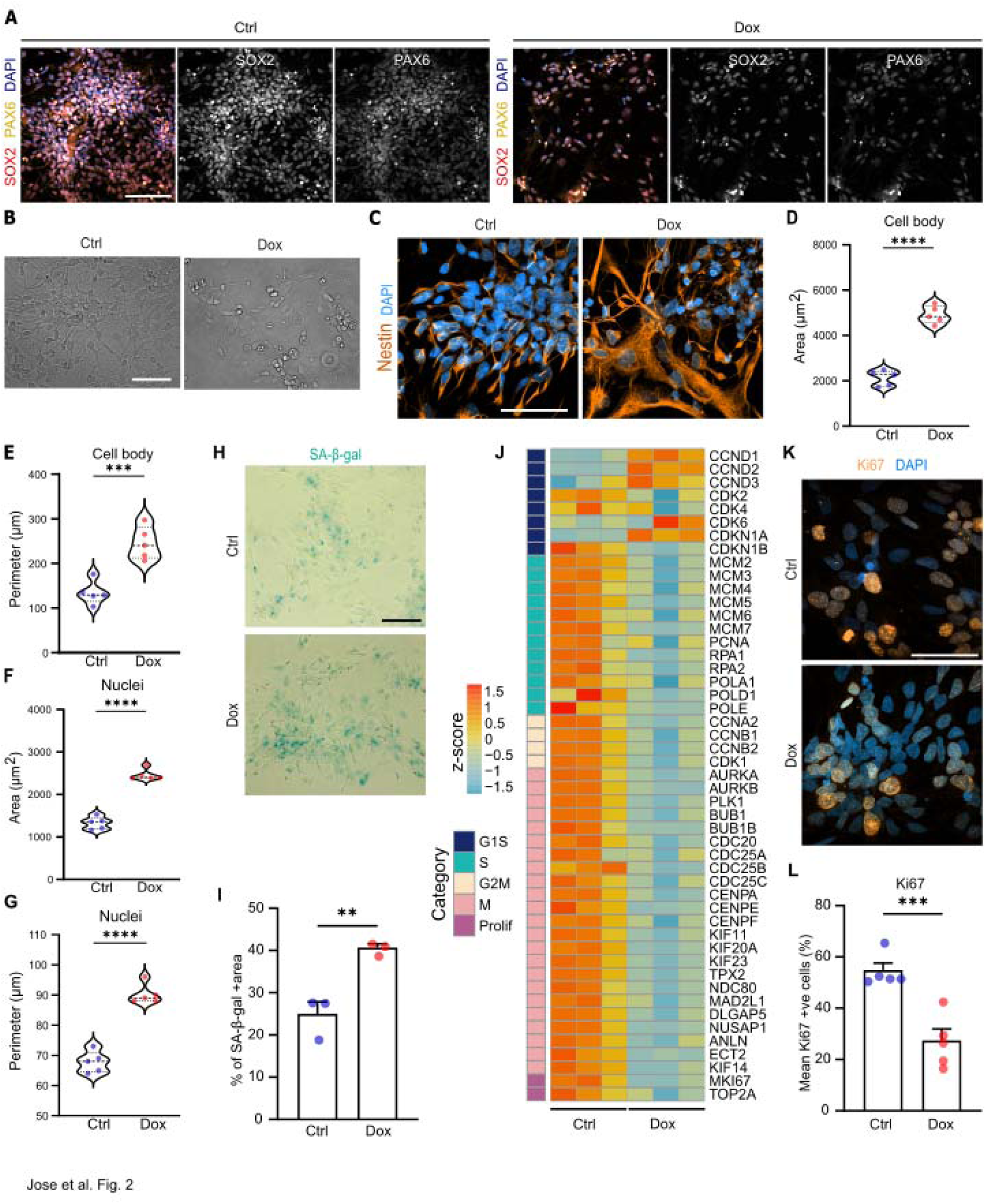
Doxorubicin induces senescence like morphological remodeling and coordinated cell-cycle suppression in NPCs. **(A)** Representative images of SOX2^+^ and PAX6^+^ cells in Ctrl and Dox-treated NPCs. **(B)** Representative brightfield images showing morphology of Ctrl and Dox-treated NPCs in culture. **(C)** Representative images of Nestin^+^ cells in Ctrl and Dox-treated NPCs. **(D-E)** Quantification of cytoplasmic areas and perimeters in Ctrl and Dox-treated NPCs. Values represent mean + SEM from *n* = 5 biological replicates, \*\*\**p* < 0.001; \*\*\*\**p* < 0.0001, unpaired two-tailed Student’s *t* test. **(F-G)** Quantification of nuclear areas and perimeters in Ctrl and Dox-treated NPCs. Values represent mean + SEM from *n* = 5 biological replicates, \*\**p* < 0.01; \*\*\*\**p* < 0.0001, unpaired two-tailed Student’s *t* test. **(H)** Representative images of senescence-associated β-galactosidase in Ctrl and Dox-treated NPCs. **(I)** Quantification of the area percentage of senescence-associated β-galactosidase staining in Ctrl and Dox-treated NPCs. Values represent mean + SEM from *n* = 3 biological replicates, \*\**p* < 0.0,1 unpaired two-tailed Student’s *t* test. **(J)** Heatmaps showing the expression patterns of cell-cycle-related genes, grouped by their respective cell-cycle stages. Each column represents an individual sample (n=3 Ctrl and n= 3 Dox-treated NPCs). Heatmaps were generated from VST-transformed, normalized gene expression values, and colors indicate the z-score of each gene across samples. **(L)** Representative images of Ki67^+^ proliferating cells in Ctrl and Dox-treated NPCs. **(M)** Quantification of the percentage of Ki67+ cells in Ctrl and Dox-treated NPCs. Values represent mean + SEM from *n* = 5 biological replicates, \*\*\**p* < 0.001, unpaired two-tailed Student’s *t* test.

To further confirm senescence induction, we assessed senescence-associated β-galactosidase (SA-β-gal) activity. Dox-treated NPCs showed a pronounced increase in SA-β-gal staining compared with controls (**Fig. 2H**), and quantitative analysis revealed a significant elevation in SA-β-gal-positive area per cell (**Fig. 2I**).

At the transcriptional level, the landscape of cell cycle regulators further supported a senescence-like state in Dox-treated NPCs. Heatmap analysis demonstrated broad suppression of proliferative and cell cycle-driving gene programs, including downregulation of markers of cell proliferation (*MKI67*, *TOP2A*), S phase (*MCM2–7*, *RPA1/2*, *POLA1*, *POLD1*, *POLE*), G2/M transition (*CCNA2*, *CCNB1/2*, *CDK1*), and mitosis (multiple *KIF* family members, *AURKA/B*, *CENPA/E/F*, *PLK1*). In contrast, genes associated with G1-S control and cell cycle-arrest including *CCND1–3*, *CDK6*, and the cyclin dependent kinase inhibitor *CDKN1A* (p21) were upregulated (**Fig. 2J**). This coordinated repression of S-, G2-, and M-phase gene expression, together with induction of G1-associated regulators and p21, is consistent with a stable G1 arrest, a hallmark transcriptional signature of senescence (Li et al. 2026). Consistently, immunofluorescence staining for Ki67 confirmed a significant reduction in the proportion of actively proliferating NPCs in the Dox-treated group compared to controls (**Fig. 2K, L**).

Together, these findings demonstrate that Dox-treated NPCs undergo coordinated morphological remodeling and cell cycle suppression, consistent with the establishment of a stable senescence-like state.

### 2.3 Doxorubicin remodels DNA damage response pathways and induces DNA damage in NPCs

To examine how Dox reshapes DDR pathways, we profiled expression of genes spanning major DDR modules, including damage sensors, homologous recombination (HR), non-homologous end joining (NHEJ), base excision repair (BER), nucleotide excision repair (NER), mismatch repair (MMR), and p53-associated stress response genes (**Fig. 3A**). Across the dataset, upstream damage sensor genes (*ATM*, *ATR*, *CHEK1, CHEK)* show comparable expression levels following Dox treatment, suggesting that early DDR checkpoint activation may occur primarily at the post-translational level rather than through transcriptional induction. In contrast, Dox treatment led to pronounced suppression of core HR components including *BRCA1*, *BRCA2*, *RAD51* and its paralogs (*RAD51B/C/D*, *XRCC2/3*), *PALB2*, and *BLM*. This is consistent with downregulation of high-fidelity repair pathways as cells exit the cell cycle during senescence. Other repair pathways exhibited more heterogeneous responses: genes involved in NHEJ, BER, NER, and MMR showed mixed moderate changes, suggesting partial but nonuniform engagement of alternative repair mechanisms. These transcriptional alterations were accompanied by increased expression of p53-responsive genes (*CDKN1A*, *GADD45A*, *BBC3*), consistent with robust p53-mediated stress signaling and sustained cell cycle arrest (**Fig. 3A**).

**Figure 3.**
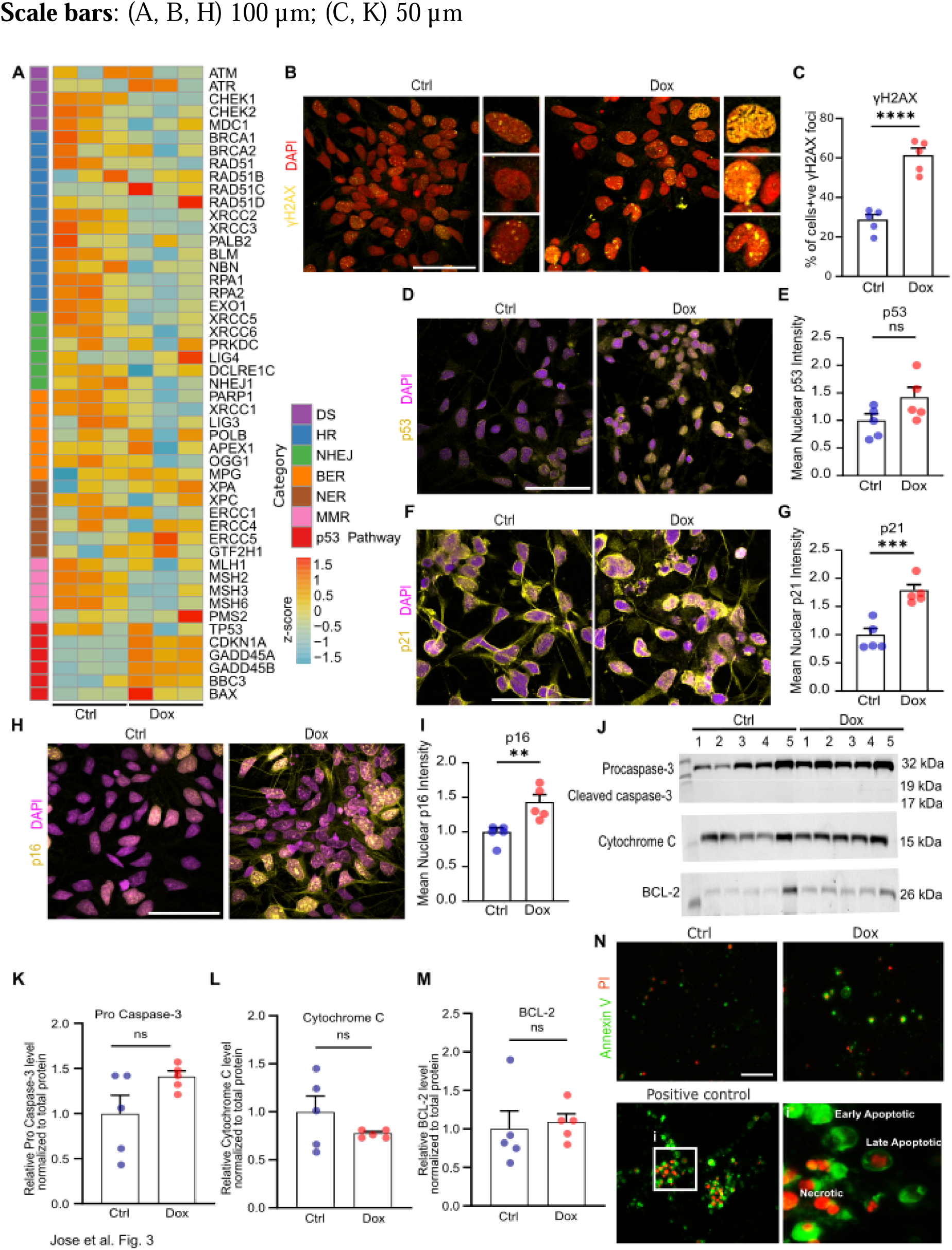
Doxorubicin remodels DNA damage response pathways and induces DNA damage in NPCs. **(A)** Heatmaps showing the expression patterns of DNA damage response (DDR) pathway related genes, grouped by major DDR categories. Each column represents an individual sample (n=3 Ctrl and n= 3 Dox-treated NPCs). Heatmaps were generated from VST-transformed, normalized gene-expression values, and colors indicate the z-score of each gene across samples. **(B)** Representative images of γH2AX^+^ foci in Ctrl and Dox-treated NPCs. **(C)** Quantification of the percentage of nuclei with γH2AX+ foci in Ctrl and Dox-treated NPCs. Values represent mean + SEM from *n* = 5 biological replicates, \*\*\*\**p* < 0.0001, unpaired two-tailed Student’s *t* test. **(D)** Representative images of p53immunofluorescence staining in Ctrl and Dox-treated NPCs. **(E)** Quantification of mean nuclear p53 fluorescence intensity in Ctrl and Dox-treated NPCs. Values represent mean + SEM from *n* = 5 biological replicates, ns; not significant, unpaired two-tailed Student’s *t* test. **(F)** Representative images of p21 immunofluorescence staining in Ctrl and Dox-treated NPCs. **(G)** Quantification of mean nuclear p21 fluorescence intensity in Ctrl and Dox-treated NPCs. Values represent mean + SEM from *n* = 5 biological replicates, \*\*\**p* < 0.001, unpaired two-tailed Student’s *t* test. **(H)** Representative images of p16 immunofluorescence staining in Ctrl and Dox-treated NPCs. **(I)** Quantification of mean nuclear p16 fluorescence intensity in Ctrl and Dox-treated NPCs. Values represent mean + SEM from *n* = 5 biological replicates, \*\**p* < 0.01, unpaired two-tailed Student’s *t* test. **(J)** Representative western blots showing the expression of apoptosis related proteins procaspase-3, cleaved caspase-3, cytochrome c and BCL-2 in Ctrl and Dox-treated NPCs. Shown blots are representative of n=5 biological replicates. **(K-M)** Quantification of procaspase-3, cytochrome c and BCL-2 protein levels in Ctrl and Dox-treated NPCs, normalized to total protein. Values represent mean + SEM from *n* = 5 biological replicates, ns; not significant, unpaired two-tailed Student’s *t* test. **(N)** Representative images of Annexin V^+^ (green) and Propidium iodide (PI)^+^ (red) staining in Ctrl and Dox-treated NPCs. A positive control (i) to show early apoptotic (Annexin V^+^/PI^−^) necrotic (Annexin V^−^/PI^+^) and late apoptotic (Annexin V^+^/PI^+^) cell populations. **Scale bars**: (B, D, F, H) 50 µm; (N) 100 µm

To directly assess DNA damage induced by Dox, we quantified γH2AX, a marker of DNA double-strand breaks. Dox-treated NPCs displayed a pronounced increase in γH2AX nuclear foci compared to controls (**Fig. 3B**), and quantitative analysis confirmed a significant increase in γH2AX-positive nuclei (**Fig. 3C**). This increase in genotoxic stress is consistent with the reduced proliferative capacity and transcriptional suppression of cell cycle regulators in Dox-treated NPCs.

Together, these findings demonstrate that Dox treatment drives extensive remodeling of DNA damage response pathways, characterized by suppression of high-fidelity repair mechanisms and accumulation of DNA damage in NPCs.

### 2.4 Transcription factor analysis reveals coordinated repression of proliferative programs and activation of stress response signaling

To elucidate the regulatory architecture underlying the transcriptional changes in Dox-treated NPCs, we performed ChIP-X–based TF enrichment analysis on both downregulated and upregulated gene sets. The downregulated genes showed strong enrichment for targets of the E2F transcription factors, particularly E2F4, E2F6, and E2F1, along with NFY (NFYA/NFYB), FOXM1, and the MYC/MAX network (**Fig. S2A**). These TFs normally drive expression of genes required for DNA replication, S-phase progression, G2/M transition, and mitosis, and their enrichment reflects the broad shutdown of cell cycle and proliferative programs during senescence (Fischer et al. 2022, Herr et al. 2024). Enrichment of BRCA1- and SIN3A-bound promoters further supports the coordinated repression of homologous recombination repair genes and chromatin-mediated silencing of cell cycle-related targets.

In contrast, the upregulated genes were strongly enriched for TP53 targets, including well-established p53-responsive stress genes such as *BBC3 (PUMA)*, *RRM2B*, *TP53INP1*, *SESN2*, *DDB2*, *ZMAT3*, and *PLK2/3*, highlighting activation of the p53-driven DDR (**Fig. S2B**). Additional enrichment for SUZ12/PRC2, SMAD4, and NFE2L2 (NRF2) indicated induction of pathways involved in chromatin remodeling, TGFβ signaling, and oxidative stress defense, respectively. Stress responsive TFs such as FOSL2 (AP-1), GATA2, and FOXA1 were also represented, reflecting transcriptional programs associated with cytoskeletal remodeling, ECM changes, and cellular stress adaptation. Together, these TF signatures reveal a clear regulatory shift from proliferation-driven transcription toward DDR, stress adaptation, and senescence-associated programs following Dox exposure.

### 2.5 p53-dependent signaling and p21/p16 induction establish stable senescence in NPCs

To further contextualize the transcriptional shifts accompanying senescence, we compared our transcriptomic data with a curated set of TP53-regulated genes defined by *Tang et al. (Tang et al. 2007)*. Among the TP53-upregulated genes, *CDKN1A*, encoding p21, showed the strongest induction in Dox-treated NPCs, alongside increased expression of additional p53-responsive targets such as *ZMAT3* and *USF2*, consistent with persistent activation of the DDR. Upregulation of *A2M*, *WFDC1*, *ABCA8*, *LCAT*, and *VAMP8* further indicated early engagement of SASP-associated and metabolic remodeling pathways. Dox exposure also increased *CCND2* and *TFAP4*, genes linked to p53-mediated modulation of cell cycle arrest (Fischer and Sammons 2024, Stracker 2024) **(Fig. S3A)**.

Conversely, TP53-downregulated genes from the *Tang et al*. dataset were broadly repressed in Dox-treated NPCs and included key regulators of cell cycle progression, mitosis, DNA replication, and chromatin organization. These included *LMNB1*, *EZH2*, *CDK1*, *CCNB2*, *TOP2A*, *BUB1*, multiple *KIF* family members, and replication licensing factors, reflecting a robust shutdown of proliferative transcriptional programs **(Fig. S3B)**.

Additionally, qPCR analysis also demonstrated significant upregulation of *CDKN1A* (encoding p21) and *CDKN2A* (encoding p16) transcripts in Dox-treated NPCs, while *TP53* (encoding p53) mRNA remained unchanged (**Fig. S4A-C**). Immunofluorescence confirmed this pattern, where p53 protein abundance was unaffected by Dox, whereas both p21 and p16 displayed significant increase (**Fig. 3D-I**). These data indicate that Dox-treatment induced cell cycle arrest in the absence of transcriptional or protein-level upregulation of p53, suggesting engagement of senescence-associated cell cycle control mechanisms operating downstream of canonical TP53 activation.

To determine whether Dox-treated NPCs had entered a stable senescent state, we extended the experimental timeline and collected cells seven days after treatment withdrawal (**Fig. S4D**). Quantitative PCR analysis revealed a sustained and pronounced upregulation of *CDKN1A* (p21) and *CDKN2A* (p16) mRNA levels at this late time point (**Fig. S4E, F)**, indicating the persistence of senescence associated cell cycle arrest and confirming that Dox treatment induced a stable senescence program in NPCs.

Together, these findings demonstrate that Dox treatment activates p53-dependent signaling and induces sustained p21 and p16 expression, establishing a stable senescence program in NPCs.

### 2.6 Doxorubicin primes apoptotic pathways but promotes apoptosis resistance in NPCs

We next assessed whether the optimized concentration of Dox used to induce senescence triggered apoptosis. Transcriptional profiling showed coordinated upregulation of upstream components of both the extrinsic apoptosis pathway (*CASP8*, *CASP10*, *FADD*, *TNFRSF10A/B*) and the intrinsic mitochondrial pathway (*BAX*, *BAK1*, *CYCS*, *DIABLO*) (**Fig. S5A**). This pattern indicates broad engagement of apoptotic signaling cascades downstream of p53. Despite this priming, executioner caspases (*CASP3*, *CASP6*, *CASP7*) showed minimal transcriptional activation, while several anti-apoptotic BCL2 family members exhibited stable or slightly increased expression. This pattern, with activation of upstream apoptotic signaling but limited executioner caspase activity, is characteristic of senescent cells. Dox also induced robust upregulation of ER-stress/UPR markers (*ATF4*, *DDIT3*, *EDEM1*, *ERN1*), supporting the presence of proteotoxic stress and metabolic strain commonly associated with the senescence program (**Fig. S5A**). These RNA-seq data indicate that Dox-treated NPCs adopt a transcriptional profile consistent with p53-driven senescence, characterized by activation of apoptotic signaling, ER stress responses, and suppression of downstream caspase activation.

Consistent with this, protein analysis revealed no significant changes in full length or cleaved caspase3 (**Fig. 3J, K**), cytochrome c (**Fig. 3J, L**), or BCL2 (**Fig. 3J, M**), confirming that Dox does not trigger mitochondrial or caspase-dependent apoptosis. Annexin V/PI staining showed only a modest increase in apoptotic populations in Dox-treated NPCs compared to control, with the majority of Dox-treated NPCs remaining viable (**Fig. 3N, Fig. S5B, C**). These findings indicate that while a small proportion of cells may undergo apoptosis or secondary necrosis, the predominant outcome of Dox exposure leads to stable entry into senescence rather than activation of the apoptotic cascade.

Collectively, functional and transcriptional analyses demonstrate that Dox treatment drives NPCs into a robust senescence-like state characterized by apoptosis resistance and reinforcement of anti-apoptotic signaling across both intrinsic and extrinsic pathways. This integrated profile is consistent with the well-established paradigm of senescence accompanied by apoptosis resistance.

### 2.7 Senescence induces mitochondrial protein remodeling, structural alterations, and metabolic reprogramming in NPCs

To investigate how senescence alters mitochondrial metabolism, we first analyzed RNA-seq data focusing on mtDNA encoded respiratory chain subunits and nuclear-encoded glycolytic and pentose phosphate pathway (PPP) genes. Heatmaps revealed a coordinated increase in the expression of mtDNA encoded respiratory chain subunits under Dox compared with control (**Fig. 4A**). Complex I subunits (*MT-ND1/2/3/4/5*), complex III (*MT-CYB*), and complex IV (*MT-CO1/2/3*) were consistently higher in Dox-treated samples, indicating enhanced mitochondrial transcription. In contrast, key glycolytic and PPP genes including *HK1*, *PGK1*, *ENO1*, *ALDOC*, *GAPDH*, *PKM, LDHA*, and *G6PD* were broadly reduced in Dox treated cells compared with controls (**Fig. 4B**). Together, these transcriptional patterns indicate a metabolic shift toward oxidative phosphorylation coupled to suppression of glycolysis/PPP, consistent with a senescence-associated metabolic phenotype.

**Figure 4.**
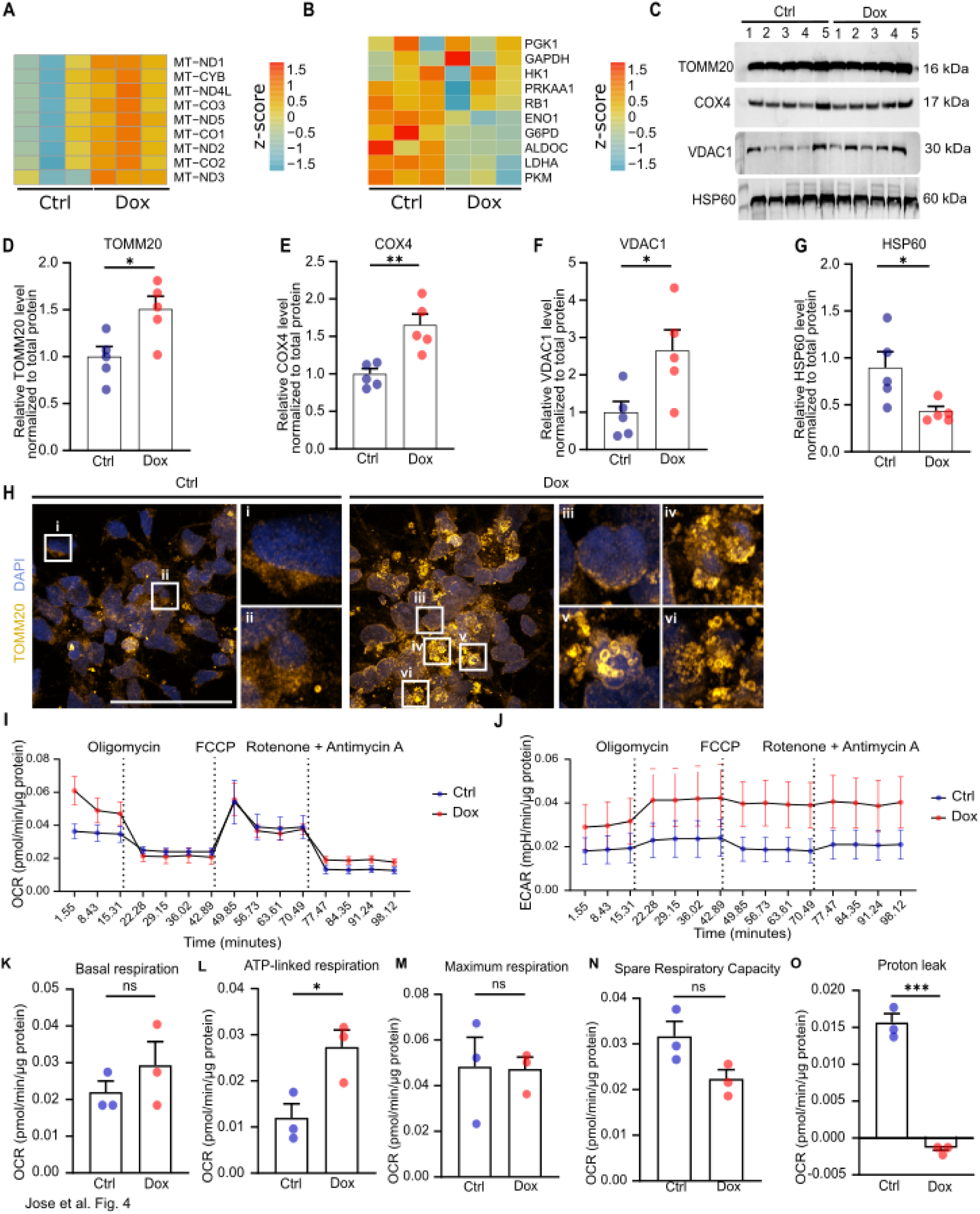
Senescence induces mitochondrial protein remodeling, structural alterations, and metabolic reprogramming in NPCs. **(A)** Heatmaps showing the expression patterns of mitochondrial respiratory chain subunits. Each column represents an individual sample (n=3 Ctrl and n= 3 Dox treated NPCs). Heatmaps were generated from VST-transformed, normalized gene-expression values, and colors indicate the z score of each gene across samples. **(B)** Heatmaps showing the expression patterns of glycolytic and pentose phosphate pathway genes. Each column represents an individual sample (n=3 Ctrl and n= 3 Dox-treated NPCs). Heatmaps were generated from VST-transformed, normalized gene expression values, and colors indicate the z-score of each gene across samples. **(C)** Representative western blots showing the expression of mitochondrial proteins TOMM20, COX4, VDAC1 and HSP60 in Ctrl and Dox-treated NPCs. Shown blots are representative of n=5 biological replicates. **(D-G)** Quantification of TOMM20, COX4, VDAC1, HSP60 protein levels in Ctrl and Dox-treated NPCs, normalized to total protein. Values represent mean + SEM from *n* = 5 biological replicates, \**p* < 0.05; \*\**p* < 0.01, unpaired two-tailed Student’s *t* test. **(H)** Representative images of TOMM20 immunofluorescence staining in Ctrl and Dox-treated NPCs. **(I)** Seahorse XF analysis of OCR (pmol/min) in Ctrl and Dox-treated NPCs, normalized to total protein content. Data represents mean + SEM from *n* = 3 biological replicates. **(J)** Seahorse XF analysis of ECAR (mpH/min) in Ctrl and Dox-treated NPCs, normalized to total protein content. Data represents mean + SEM from *n* = 3 biological replicates. **(K-N)** Quantification of basal, ATP-linked and maximum respiration, and spare respiratory capacity derived from Seahorse XF OCR measurements. Values represent mean + SEM from *n* = 3 biological replicates, normalized to total protein content, \**p* < 0.05; ns: not significant, unpaired two-tailed Student’s *t* test. **(O)** Quantification of proton leak derived from Seahorse XF ECAR measurements. Values represent mean + SEM from *n* = 3 biological replicates, normalized to total protein content,; \*\*\**p* < 0.001, unpaired two-tailed Student’s *t* test. **Scale bars**: (H) 50 µm.

To determine how senescence impacts mitochondrial structure and function in NPCs, we examined key mitochondrial proteins involved in import (TOMM20), oxidative phosphorylation (COX4), membrane permeability and metabolite exchange (VDAC1) and proteostasis (HSP60). Western blot analysis revealed increased abundance of TOMM20, COX4, and VDAC1 in Dox treated senescent cells compared with controls (**Fig. 4C-F**), while HSP60 levels were significantly reduced, consistent with impaired mitochondrial chaperone function (**Fig. 4G**). These protein-level changes indicate broad mitochondrial reprogramming and are consistent with previous reports that Dox-induced senescence is associated with increased mitochondrial stress and compensatory mitochondrial adaptation (Linders et al. 2024).

Dox-treated NPCs exhibited pronounced alterations in mitochondrial organization consistent with senescence-associated mitochondrial dysfunction (SAMD) (Hruby and Higuchi-Sanabria 2025, Korolchuk et al. 2017, Zhang et al. 2025). TOMM20 immunostaining revealed increased mitochondrial signal intensity, indicative of mitochondrial accumulation, a recognized feature of senescent cells in which impaired mitophagy permits the persistence of dysfunctional organelles (**Fig. 4H**). In addition, Dox-treated NPCs displayed numerous donut-shaped mitochondria, a stress-associated morphology linked to altered fusion–fission dynamics, mitochondrial swelling, and membrane-potential stress in hypoxia-reoxygenation and related conditions (Liu and Hajnoczky 2011). Mitochondria also adopted a predominantly perinuclear distribution, consistent with stress induced, dynein/microtubule dependent retrograde transport reported to increase nuclear ROS and facilitate HIF-dependent transcription (Agarwal and Ganesh 2020, Al-Mehdi et al. 2012) (**Fig. 4H**).

We next assessed mitochondrial function using Seahorse extracellular flux analysis (**Fig. 4I, J**). Basal respiration was not significantly different between conditions, indicating that steady-state mitochondrial activity is preserved in Dox-treated NPCs (**Fig. 4K**). In contrast, Dox-treated NPCs displayed significantly increased ATP-linked respiration compared to controls, indicating enhanced coupling of mitochondrial respiration to ATP production (**Fig. 4L**). Maximal respiration was also unchanged (**Fig. 4M**), suggesting that overall respiratory capacity remains intact. However, Dox-treated NPCs showed a trend toward reduced spare respiratory capacity, consistent with a diminished ability to respond to increased energetic demand (**Fig. 4N**). In addition, proton leak was significantly reduced, indicating decreased mitochondrial uncoupling (**Fig. 4O**). Together, these findings indicate that senescence promotes a more constrained oxidative metabolic phenotype, characterized by preserved respiratory capacity but increased coupling to ATP production and reduced metabolic flexibility. These findings are consistent with a senescence-associated metabolic rewiring in which NPCs shift from a glycolytic to a more oxidative, tightly coupled state.

Together, transcriptomic, proteomic, morphological, and bioenergetic analyses indicate that Dox-induced senescence in NPCs is associated with mitochondrial remodeling, characterized by increased respiratory coupling, altered mitochondrial organization, and reduced metabolic flexibility, alongside suppression of glycolytic and pentose phosphate pathways. This profile is consistent with a senescence-associated shift toward a constrained oxidative metabolic state, rather than a simple increase in overall respiratory capacity.

### 2.8 Doxorubicin-induced senescence activates a pro-angiogenic SASP and TGF**β**–ECM remodeling program in NPCs

Given that the SASP is a hallmark of cellular senescence, we assessed whether Dox-treated NPCs acquire a SASP. RNA sequencing revealed a robust SASP signature, characterized by upregulation of proinflammatory cytokines and chemokines (*IL1A*, *IL1B*, *IL11*, *GDF15*, *CCL2*, *CCL20*, *CXCL1/2/3*), adhesion and immune modulators (*ICAM1*), and TGFβ ligands (*TGFB1*, *TGFB3*) in Dox-treated NPCs (**Fig. 5A**). These changes are consistent with therapy-induced senescence, in which persistent DNA damage signaling and chromatin remodeling activate NFκB/C/EBP-driven SASP networks and paracrine tissue remodeling. qPCR confirmed increased *IL1A*, *CCL2*, and *TGFB1* transcripts (**Fig. 5B-D**).

**Figure 5.**
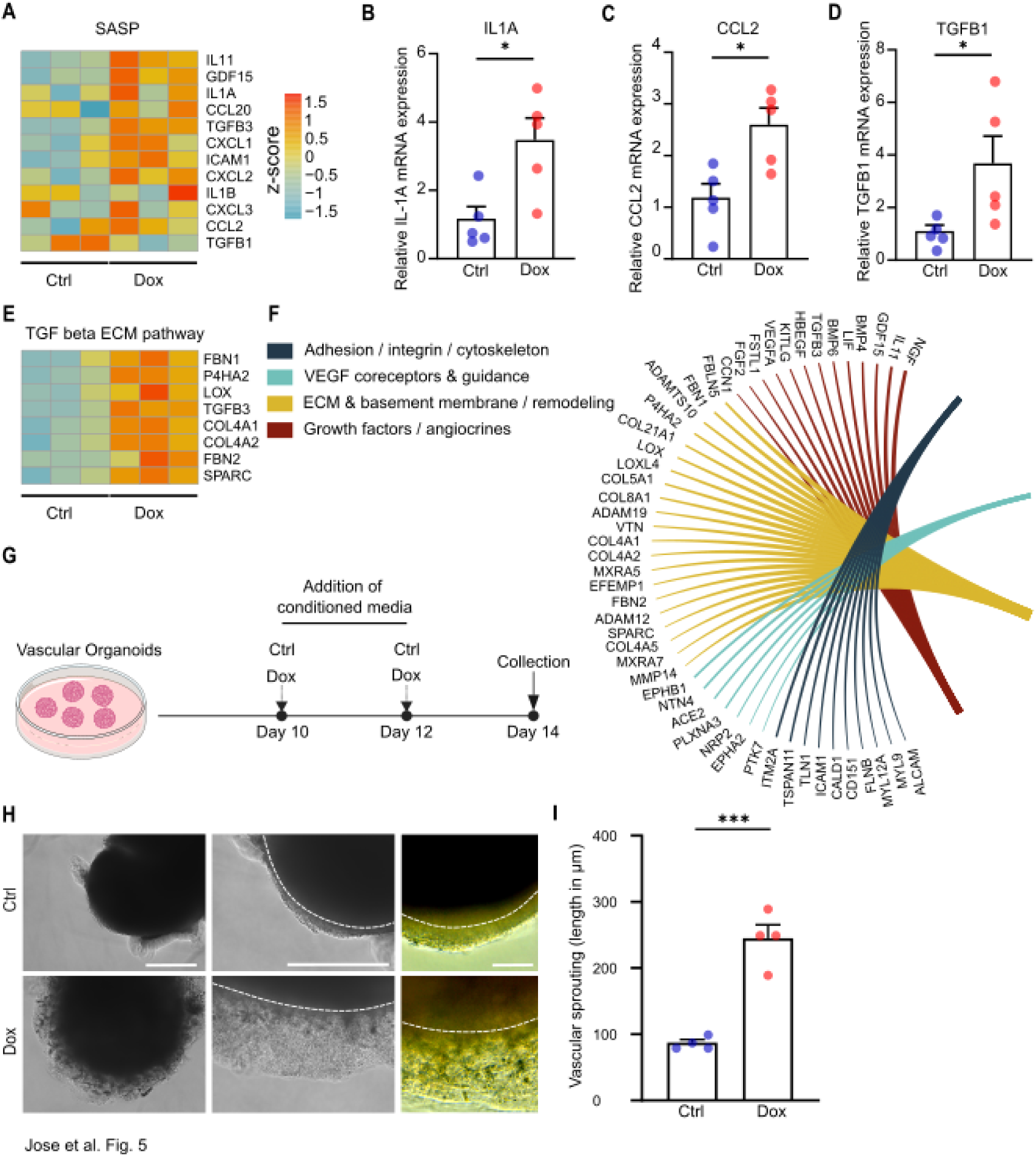
Doxorubicin induces SASP, TGFβ–ECM remodeling and proangiogenic programs that drive paracrine angiogenesis. **(A)** Heatmaps showing the expression patterns of senescence-associated secretory phenotype (SASP) related genes. Each column represents an individual sample (n=3 Ctrl and n= 3 Dox-treated NPCs). Heatmaps were generated from VST-transformed, normalized gene-expression values, and colors indicate the z-score of each gene across samples. **(B-D)** Relative mRNA expression levels of *IL1A*, *CCL2*, and *TGFB1* in Ctrl and Dox-treated NPCs measured by RT–qPCR. Values represent mean + SEM from n = 5 biological replicates, normalized to the housekeeping genes *RPS18* and *18S*, \**p* < 0.05, unpaired two-tailed Student’s *t* test. **(E)** Heatmaps showing the expression patterns of TGF beta and extracellular matrix (ECM) pathway related genes. Each column represents an individual sample (n=3 Ctrl and n= 3 Dox-treated NPCs). Heatmaps were generated from VST-transformed, normalized gene-expression values, and colors indicate the z-score of each gene across samples. **(F)** Radial plot showing the functional categorization of genes associated with adhesion/integrin/cytoskeleton organization, VEGF coreceptors and guidance, ECM and basement-membrane remodeling and growth factors/angiocrines signaling. Genes are grouped according to pathway assignment, with colored segments indicating each category. **(G)**Schematic of the experimental workflow. Conditioned medium from Ctrl and Dox-treated NPCs (collected at day 6) was applied to day 10 vascular organoids every other day. Organoids were assessed for vascular sprouting on day 14. **(H)**Representative brightfield images of vascular organoids showing vascular sprouting following 4-days exposure to conditioned medium from Ctrl and Dox-treated NPCs. **(I)**Quantification of length of vascular sprouting following exposure to conditioned medium from Ctrl and Dox-treated NPCs. Values represent mean + SEM 3-4 images from 4 independent vascular organoids, \*\*\**p* < 0.001, unpaired two-tailed Student’s *t* test. **Scale bars:** (H) 500 µm (left and middle panels), 100 µm (right panels).

Transcriptional signatures further indicated activation of a TGFβ–ECM program with over-expression of matrix/cross-linking genes (*FBN1*, *FBN2*, *LOX*, *P4HA2*), basement membrane-collagens (*COL4A1*, *COL4A2*), and the matricellular factor *SPARC* (**Fig. 5E**), coherent with TGFβ’s role in ECM deposition, cross-linking, and tissue stiffness in senescence contexts (Tominaga and Suzuki 2019). In parallel, multiple adhesion, integrin, cytoskeletal, VEGF, co-receptor/guidance, and ECM remodeling genes were upregulated, indicating a shift toward pro-migratory, pro-angiogenic signaling (**Fig. 5F**). Aligned with stress-responsive mitochondrial relocalization pathways that enhance nuclear ROS production and HIF-1-dependent *VEGF* transcription (Al-Mehdi et al. 2012), our RNA sequencing data showed upregulation of *VEGFA* in Dox-treated-NPCs (**Fig. 5F**), suggesting a mechanistic link between mitochondrial stress, SASP/TGFβ remodeling, and pro-angiogenic output.

To functionally validate these findings, human iPSC-derived vascular organoids were exposed to conditioned media from control and Dox-treated NPCs culture (**Fig. 5G**). Vascular organoids treated with conditioned media from Dox-treated NPCs exhibited significantly greater angiogenesis than controls (**Fig. 5H, I**), indicating that senescent NPC secretome comprising inflammatory cytokines/chemokines, TGFβ-dependent ECM modulators, and VEGF axis cues are sufficient to drive endothelial growth and network expansion in vitro.

Together, these findings demonstrate that Dox-induced senescence reprograms NPCs into a pro-angiogenic SASP that drives vascular remodeling through paracrine signaling.

### 2.9 Doxorubicin induces a stable senescent state in NPCs in hippocampal organoids

In monolayer NPCs, Dox treatment established a stable senescent state characterized by (i) a senescence shifted transcriptional landscape, (ii) altered cell morphology and reduced proliferation, (iii) activation of the DNA-damage response with p53/p21/p16 induction, (iv) mitochondrial/energy-metabolism rewiring, and (v) a SASP with pro-angiogenic secretome features.

To determine whether Dox-induced senescence extends to a tissue-like environment, we leveraged human iPSC-derived hippocampal organoids (Soper et al. 2025). Unlike 2D cultures, hippocampal organoids recapitulate key aspects of brain architecture, including cellular heterogeneity and spatial organization of stem cell niche, enabling NPCs to interact with surrounding neuronal and glial populations (**Fig. 6A**). This system therefore provides a platform to assess senescence within a complex microenvironment that more closely mimics the in vivo hippocampal niche. Organoids were treated with Dox to evaluate whether senescence-associated phenotypes are similarly induced in this 3D context (**Fig. 6B**). SA-β-gal staining supported senescence induction within NPC layers with increased staining observed in Dox-treated organoids (**Fig. 6C**). Furthermore, Dox-treated organoids exhibited a reduction in SOX2^+^ NPCs (**Fig. 6D, E**). Consistently, immunofluorescence analysis for proliferating (Ki67^+^) and mitotic (pH3^+^) NPCs revealed a significant decrease in both populations following Dox treatment, from 17.8% to 3.9% and 1.5% to 0.03%, respectively (**Fig. 6D, F-H**). This reduction in proliferative activity coincided with enhanced NPC DNA damage, as indicated by elevated numbers of γH2AX foci within SOX2^+^ nuclei, with 54% of SOX2^+^ nuclei in Dox-treated organoids displaying 1-3 γH2AX foci, compared to just 7% in control organoids (**Fig. 6I-K**). Additionally, the expression of both p53 and p21 were upregulated in Dox-treated NPCs, supporting DNA-damage response activation with p53 and p21 induction (**Fig. 6L-O**).

**Figure 6:**
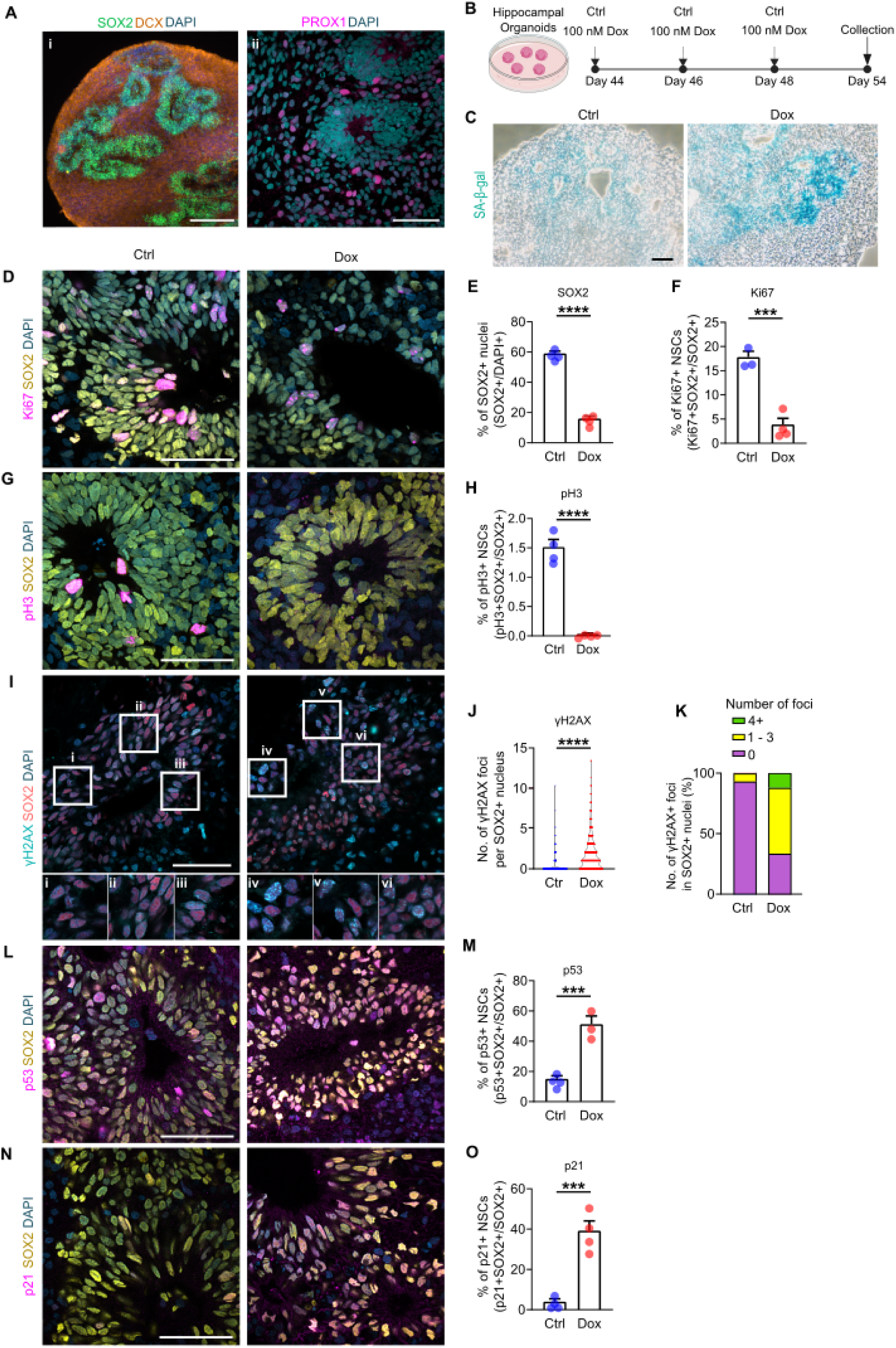
Doxorubicin induces a stable senescent state in NPCs in hippocampal organoids. **(A)** Representative images of hippocampal organoids showing SOX2+ stem cell layer and DCX^+^ immature neurons (i) and PROX1^+^ dentate granule neurons (ii) at day 50 of culture. **(B)** Schematic of the experimental workflow. Hippocampal organoids were treated with 100 nM Dox, or vehicle control (Ctrl), on days 44, 46 and 48 of culture. Organoids were harvested on day 54 of culture for downstream analysis. **(C)** Representative images of senescence-associated β-galactosidase in Ctrl and Dox-treated hippocampal organoids. **(D)**Representative images of Ki67^+^ proliferating SOX2^+^ NPCs in Ctrl and Dox-treated hippocampal organoids. **(E)**Quantification of the proportion of SOX2^+^ NPCs in Ctrl and Dox-treated hippocampal organoids. N = 23 - 24 sections from 4 independent hippocampal organoids per condition, \*\*\*\**p* < 0.0001, unpaired two-tailed Student’s *t* test. **(F)**Quantification of the proportion of Ki67^+^ proliferating SOX2^+^ NPCs in Ctrl and Dox-treated hippocampal organoids. N = 11 - 15 sections from 3 - 4 independent hippocampal organoids per condition, \*\*\**p* < 0.001, unpaired two-tailed Student’s *t* test. **(G)** Representative images of pH3^+^ mitotic SOX2^+^ NPCs in Ctrl and Dox-treated hippocampal organoids. **(H)** Quantification of the proportion of pH3^+^ mitotic SOX2^+^ NPCs in Ctrl and Dox-treated hippocampal organoids. N = 10 sections from 4 independent hippocampal organoids per condition, \*\*\*\**p* < 0.0001, unpaired two-tailed Student’s *t* test. **(I)** Representative images of γH2AX^+^ foci in SOX2^+^ NPCs in Ctrl and Dox-treated hippocampal organoids. **(J)** Quantification of the total number of γH2AX^+^ foci in SOX2^+^ NPCs in Ctrl and Dox-treated hippocampal organoids. N = 1507 – 2109 nuclei from 9 sections from 3 independent hippocampal organoids per condition, \*\*\*\**p* < 0.0001, Mann-Whitney U test. **(K)** Quantification of the number of γH2AX^+^ foci per nucleus in SOX2^+^ NPCs in Ctrl and Dox-treated hippocampal organoids. N = 1507-2109 nuclei from 9 sections from 3 independent hippocampal organoids per condition. **(L)** Representative images of p53 immunofluorescence staining in SOX2^+^ NPCs in Ctrl and Dox-treated hippocampal organoids. **(M)** Quantification of the proportion of p53^+^ SOX2^+^ NPCs in Ctrl and Dox-treated hippocampal organoids. N = 9 sections from 3 - 4 independent hippocampal organoids per condition, \*\*\**p* < 0.001, unpaired two-tailed Student’s *t* test. **(N)** Representative images of p21 immunofluorescence staining in SOX2^+^ NPCs in Ctrl and Dox-treated hippocampal organoids. **(O)**Quantification of the proportion of p21^+^ SOX2^+^ NPCs in Ctrl and Dox-treated hippocampal organoids. N = 13 – 14 sections from 4 independent hippocampal organoids per condition, \*\*\**p* < 0.001, unpaired two-tailed Student’s *t* test. **Scale bars:** (A) (i) 200 µm (ii) 50 µm; (C) 100 µm; (D, G, I, L, N) 50 µm.

Together, these data show that Dox consistently induces senescence in NPCs across both 2D and 3D culture systems, including NPC monolayers and 3D hippocampal organoids. This establishes a tractable human model to interrogate cellular and molecular mechanisms of senescence relevant to hippocampal development, neurogenesis, as well as aging, and neurodegeneration-associated tissue remodeling.

## 3. Discussion

Aging is the strongest risk factor for neurodegenerative disorders, and a progressive decline in the brain’s regenerative capacity is increasingly recognized as a key contributor to this vulnerability. Reduced hippocampal neurogenesis has been associated with cognitive impairment in aging and neurodegenerative disease, highlighting the importance of understanding how NPC function is compromised in these contexts. In this study, we establish a human model of DNA damage–induced senescence in NPCs, providing a tractable system to investigate how senescence-associated changes may contribute to impaired progenitor function during brain aging and disease.

In agreement with previous reports showing diminished NPC abundance and reduced immature neuronal populations in the aging human hippocampus (Boldrini et al. 2018, Franjic et al. 2022, Moreno-Jimenez et al. 2019, Tobin et al. 2019), our findings suggest that cellular senescence may contribute to stem cell decline following genotoxic stress. This is particularly relevant given that preservation of neurogenic markers in older individuals strongly correlates with maintained cognitive function (Disouky et al. 2026, Terreros-Roncal et al. 2021), implying that even modest senescence-associated impairments could have significant consequences for hippocampal resilience.

A major challenge in the field has been distinguishing bona fide senescence phenotypes from the acute cytotoxic effects of DNA damage. By employing a transient low dose exposure to Dox, a well-characterized inducer of stress-induced premature senescence (SIPS) (Du et al. 2025, Gao et al. 2023, Singh et al. 2025), we establish a controlled human senescent NPC model that circumvents this limitation. Importantly, the senescence program observed in NPCs closely mirrors canonical signatures described in fibroblasts, epithelial cells, and other stem cell systems, supporting the concept that core senescence mechanisms are conserved across tissues. At the same time, our transcriptomic analyses reveal NPC-specific responses, including alterations in cholesterol metabolism, highlighting how cell identity shapes the manifestation and functional consequences of senescence. Consistent with the establishment of a stable senescent state, Dox-treated NPCs exhibited robust activation of p53-dependent pathways, induction of p21 and p16, repression of cell cycle and DNA replication genes, and persistent accumulation of γH2AX foci. The persistence of this response several days after drug withdrawal indicates that DDR activation drives the establishment of a stable senescent state in NPCs. Given that DDR activity increases with age in many tissues, including the brain, DDR-driven stable senescence may contribute to the progressive depletion of the progenitor pool in vivo.

Importantly, these findings were recapitulated in human iPSC-derived hippocampal organoids, which provide a three-dimensional, multicellular context that more closely reflects in vivo tissue organization. Within this system, NPC populations exhibited hallmark features of senescence, including reduced proliferation, cell cycle repression, activation of p53 signaling, p21 upregulation, and widespread γH2AX accumulation. These observations demonstrate that the senescence program is not restricted to simplified monolayer cultures but is preserved within complex, tissue-like environments. Collectively, this highlights the potential of organoid systems as a platform to investigate how microenvironmental interactions influence senescence phenotypes during brain aging and disease.

Consistent with established features of senescence, we observed that NPCs adopt an apoptosis-primed yet apoptosis-resistant state. Although components of apoptotic signaling pathways were strongly upregulated, activation of executioner caspases remained minimal, and overall cell viability was largely preserved. This “primed but protected” configuration is a hallmark of senescent cells (Childs et al. 2015) and may enable damaged progenitors to persist long term within neurogenic niches. In the hippocampal context, the accumulation of such long-lived senescent NPCs could have important functional consequences, particularly through SASP-mediated paracrine effects on neighboring stem cells, neurons, and glial populations.

Our data further indicates that mitochondrial remodeling is a central component of NPC senescence. The observed increase in mitochondrial mass and electron transport chain components, coupled with reduced expression of glycolytic genes and altered oxidative phosphorylation dynamics, is consistent with mitochondrial dysfunction-associated senescence (MiDAS). Increased reliance on mitochondrial respiration may support the biosynthetic demands of the SASP, but may also impose metabolic rigidity, limiting the capacity of NPCs to respond to physiological cues that normally promote proliferation or differentiation. Given accumulating evidence that mitochondrial dysfunction contributes to age-related declines in neurogenic potential (Hruby and Higuchi-Sanabria 2025, Somasundaram et al. 2024, Zhang et al. 2025), our findings place NPC senescence within a broader metabolic framework relevant to both healthy aging and neurodegenerative disease. Notably, mitochondrial stress has been linked to increased ROS production and activation of HIF-1-dependent transcriptional programs, providing a potential mechanistic link between metabolic reprogramming and the pro-angiogenic SASP observed in our system.

Several limitations should be considered. First, while Dox is a well-established inducer of senescence, it represents a genotoxic stress model that may not fully capture the complexity of physiological aging. Second, although organoids provide a more physiologically relevant system than 2D cultures, they lack full vascularization and immune components present in vivo. Future studies incorporating in vivo models or more complex co-culture systems will be important to validate and extend these findings.

## 4. Conclusion

Together, these results establish a robust human model of DNA damage–induced senescence in neural progenitor cells. By integrating transcriptomic, metabolic, morphological, and molecular analyses, this system recapitulates key hallmarks of senescence while preserving NPC identity and physiological relevance. The concordance between 2D and 3D systems suggests that susceptibility to DDR-mediated senescence is an intrinsic property of human neural progenitors, with potential implications for hippocampal plasticity, cognitive aging, and neurodegenerative disease. Collectively, this integrated framework provides a platform to investigate both cell-intrinsic and niche-level consequences of senescence in the human hippocampus and may support future efforts to develop strategies aimed at preserving or restoring neurogenic capacity.

## 5. Methods and Materials

### 5.1 NPC 2D culture

Human iPSC-derived Neural Progenitor Cells (Stemcell Technologies, #200-0620) were grown in Neural Progenitor Medium (Stemcell Technologies, #05834) with Neural Progenitor supplements (Stemcell Technologies, #05836; #05837) at 37°C in a humidified 5% CO_2_ atmosphere, according to the manufacturer’s protocol (Stemcell Technologies).

### 5.2 Culture of human hippocampal organoids

Hippocampal organoids were generated from human iPSCs (Kolf2.1J) using a stepwise neural induction and regional specification protocol (Soper et al. 2025). iPSCs were maintained on Matrigel in mTeSR Plus (Stemcell Technologies, #100-0276) at 37 °C and 5% CO_2_. At ∼80% confluence, cells were dissociated and seeded as embryoid bodies (EBs) at 3 × 10^4^ cells/well in ultra-low attachment 96-well plates with 10 µM Y-27632, followed by centrifugation (300 × g, 5 min, 4 °C).

From day 2, EBs were transferred to neural induction medium with Y-27632 for 48 h, then cultured on an orbital shaker (120 rpm) with media changes every 48 h. Neural induction (days 2–5; hippocampal organoids media (HO) 1) was performed using dual SMAD and WNT inhibition. Organoids were then sequentially patterned using defined media: HO2 (days 8–9) with CHIR99021 and BMP7, followed by HO3 (days 10–19) with reduced CHIR99021 and BMP7 supplementation.

From day 20–25, organoids were gradually transitioned to maturation medium (HO4). From day 26 onwards, organoids were maintained in HO4 containing BDNF, dibutyryl cAMP, and ascorbic acid to support neuronal differentiation and maturation, with continued shaking and media changes every 48 h.

### 5.3 Generation of human blood vessel organoids

Human iPSC-derived vascular organoids (VOs) were generated as previously described (Dao et al. 2024) . On day 10, organoids were exposed to a 1:1 mixture of fresh VO medium and conditioned medium derived from control or Dox-treated NPCs. Treatments were applied twice on alternating days, and organoids were collected for analysis on day 14.

### 5.4 Senescence induction

Senescence was induced by treating cells with the chemotherapeutic agent doxorubicin (Dox). The applied dosage of Dox (Cell Signaling, #5927) as well as treatment duration were optimized for the respective senescent cell model included in this study. For genotoxic stress-induced senescence, NPCs (P2 – P6) were treated with 25 nM Dox for 24 h. Subsequently, cells were washed with Dulbecco’s phosphate-buffered saline (DPBS; Thermo Fisher Scientific) and cultured in Neural Progenitor medium for an additional 3 days. For extended culture conditions, untreated control cells were collected at day 4, whereas Dox-treated cells were maintained in drug-free medium for an additional 7 days following drug exposure. Cells were harvested at the end of incubation for further processing. Proliferating cells with no drug treatment served as controls. Cells used for the conditions were at about ∼70% confluency before treatments. Meanwhile, 100 nM Dox was administered to hippocampal organoids on days 44, 46, and 48 of culture. Vehicle controls received an equivalent volume of dH2O. Organoids were harvested for downstream analysis on day 54 of culture.

### 5.5 Senescence associated **β**-galactosidase staining

Cells were seeded at a density of 1 × 10^5^ cells/cm², and senescence-associated β-galactosidase (SA-β-Gal) activity was assessed using the Senescence β-Galactosidase Staining Kit (Cell Signaling, #9860) following the manufacturer’s instructions. Briefly, cells were fixed for 10 minutes and incubated with SAβGal staining solution (pH 6.0) for 16 hours at 37 °C in a CO_2_ free dry incubator. Images were acquired with an EVOS XL Core inverted microscope (Thermo Fischer Scientific) using a 20× objective for NPCs, and a 4× objective for hippocampal organoids.

### 5.6 Immunocytochemistry and Image Acquisition

The following antibodies were used for immunostaining: rat anti-SOX2 (1:1000, Invitrogen #14-9811-82), mouse anti-PAX6 (1:500, Proteintech #67529-1-Ig), chicken anti-Nestin (1:1000, Invitrogen #PA5-143578), rabbit anti-Ki67 (1:500, Abcam #ab15580), rabbit anti-γH2AX (1:1000, Proteintech, #29380-1-AP), rabbit anti-p53 (1:1000, Proteintech, #10442-1-AP), rabbit anti-p21 (1:500, Proteintech, #10355-1-AP), rabbit anti-p16 (1:500, Proteintech, #10883-1-AP) , rabbit anti-TOMM20 (1:1000, Abcam #ab186735). Alexa Fluor-conjugated secondary antibodies, including goat anti-rabbit IgG Alexa Fluor 488 (1:500, #A-11008), goat anti-mouse IgG Alexa Fluor 647 (1:500, #A21235), goat anti-rat IgG Alexa Fluor 488 (1:500, #), goat anti-chicken IgG Alexa Fluor 594 (1:500, #), were obtained from Thermo Fisher Scientific.

Cells were fixed in ice-cold methanol for 10 min at -20°C and subsequently rinsed three times with ice-cold Hank’s Balanced Salt solution (HBS). After fixation, cells were permeabilized and blocked in HBS solution containing 0.1% Triton-X-100 (BioXtra, #T9284), 1% goat serum (Abcam, #AB7481), 2% Bovine Serum Albumin (Sigma, #A8022) and 1% milk powder. Following cell blocking for 30 min at room temperature, cells were incubated with primary antibodies at 4°C overnight. Cells were then washed three times for 5 min each in HBS solution containing 0.1% Tween 20 and incubated with Alexa Fluor-tagged secondary antibodies for 1 h at room temperature in the dark. For antibody dilutions, a HBS solution containing 0.1% Triton-X-100, 1% goat serum and 2% Bovine Serum Albumin (BSA) was used. After three 5-min washes in HBS solution containing 0.1% Tween 20, cell nuclei were stained with 4′,6-diamidino-2-phenylindole (DAPI, 1:1000, Invitrogen #P36970) for 5 min at room temperature. After three washes in HBS solution, the cells were mounted using ProLong^TM^ Diamond Antifade Mountant (Invitrogen, #P36970).

Hippocampal organoids were fixed in 4% paraformaldehyde (PFA) for 30 minutes at room temperature followed by cryoprotection in 30% sucrose for at least 24 hours. Samples were embedded in OCT within cryomolds, frozen on dry ice, and stored at −80°C. Serial 20 μm sections were obtained for histology analysis. Samples were washed in 1X TRIS-buffered saline containing 0.05% Triton X-100 (TBS+). Antigen retrieval was performed in 1X DAKO Target Retrieval Solution. Blocking was performed using 10% BSA solution. Primary and secondary antibodies were diluted in TBS+ containing 3% BSA.

Imaging was performed on an EVOS M5000 inverted microscope (Thermo Fisher Scientific) using 10×, 20× and 40× objectives. Representative images were acquired using a Zeiss confocal microscope with 10× and 63× oil immersion objectives. Settings for gain and laser intensity were optimized for representative images. Images were processed and analyzed using CellProfiler v.4.2.6 and Fiji/ImageJ software. Quantitative image analysis parameters included cell number, nuclear and cytoplasmic intensity, cellular and nuclear area and perimeter.

### 5.7 RNA Extraction, reverse transcription and quantitative real-time PCR (qPCR)

Total RNA was isolated using RNeasy Mini Kit (Qiagen,#74536) according to the manufacturer’s protocol. cDNA was synthesized using Tetro cDNA Synthesis kit (Meridian Bioscience, #BIO-65043). qPCR was performed using GoTaq qPCR Master Mix (Promega, #A6002) on a LightCycler®480 System (Roche). Target mRNA levels were normalized to reference gene expression. The internal controls were *RPS18* and *18S*. Forward and reverse primers (all purchased from Sigma) used are listed in Table 1.

**Table 1.**
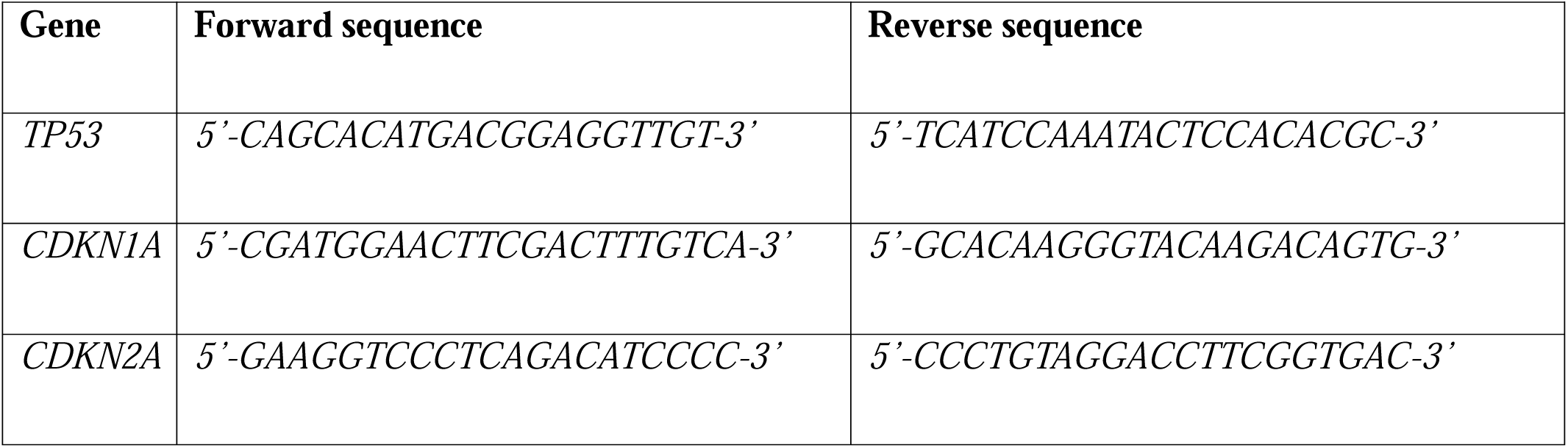
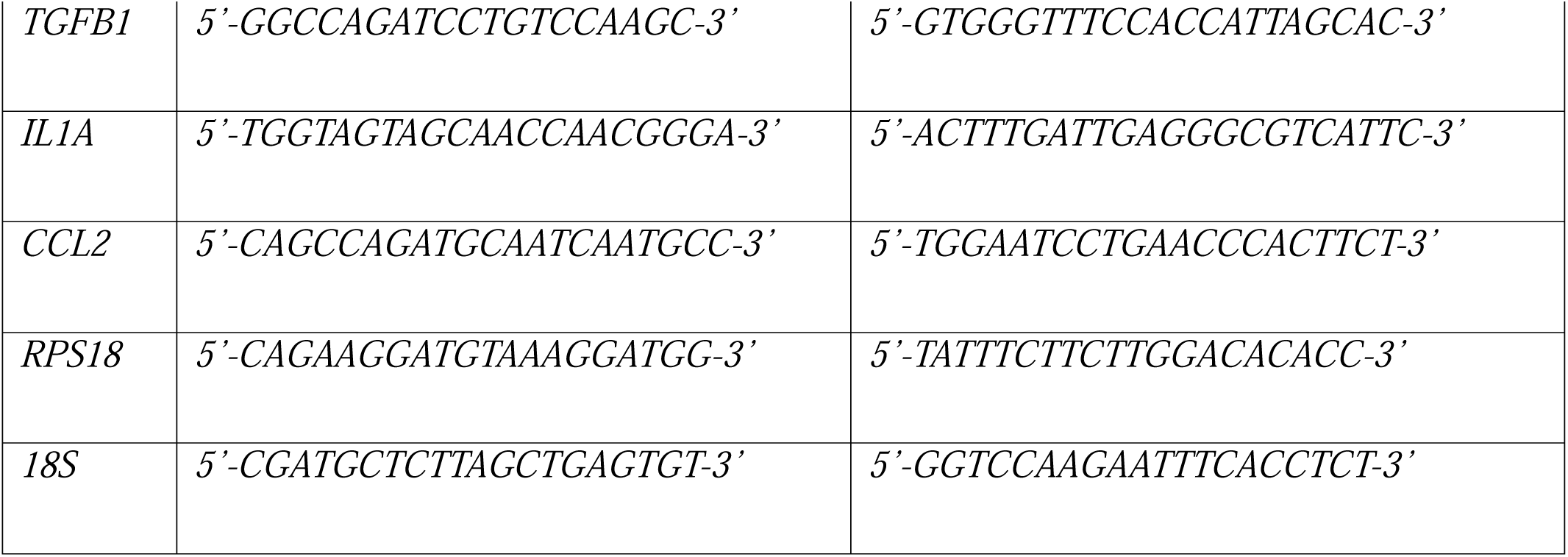
Primer sequences for real-time PCR.

### 5.8 RNA sequencing and enrichment analysis of transcriptomic data

Total RNA was isolated from cells using RNeasy Mini Kit (Qiagen, #74536) according to the manufacturer’s protocol, including on-column DNase I treatment to remove residual genomic DNA. Library preparation and paired-end sequencing was performed externally on Illumina NovaSeq X Plus Sequencing System (Novogene).

Paired-end FASTQ files were initially quality checked and trimmed using FASTP (Chen et al. 2018, Chen 2023). Processed reads were then mapped to the GRCh38 human reference genome with Salmon v1.10.2 (Patro et al. 2017) in its mapping-based mode to obtain transcript-level quantifications. Transcript- and gene-level abundance estimates were summarized using the tximeta package (Love et al. 2020) in Bioconductor, which automatically annotates quantifications by linking to reference metadata; annotations were based on Ensembl Homo sapiens release 113. Gene-level count matrices were imported into DESeq2 for differential gene expression analysis (Love, Huber and Anders 2014), applying thresholds of |log^2^ fold change| ≥ 0.4 and FDR-adjusted *p*-value < 0.05 to define differentially expressed genes. Data visualization, including volcano plots, was carried out in R using ggplot2 (version 3.5.1) in combination with ggrepel (version 0.9.6).

Functional enrichment analyses of differentially expressed genes were performed using the Enrichr web platform (Chen et al. 2013, Kuleshov et al. 2016, Xie et al. 2021). An FDR-adjusted *p*-value < 0.05 threshold was used to define functionally enriched pathways. Normalized gene-level count data were transformed using the variance-stabilizing transformation (VST) implemented in the DESeq2 package, and the resulting values were used to generate heatmaps. Heatmaps were produced using the pheatmap R package. A list of senescence-related genes was obtained from the CellAge (Avelar et al. 2020) (https://genomics.senescence.info/cells/) and SeneQuest (Gorgoulis et al. 2019) databases.

### 5.9 Western blot

Cells were lysed in RIPA buffer (1.5 M TrisHCl, 5 M NaCl, 5% SDS, 1% Triton X100, 5% sodium deoxycholate, 0.1 M EDTA, 100 mM NaF, 1 M Na_3_VO_4_; pH 7.4) supplemented with 1× Protease Inhibitor Cocktail (Roche) and 1× PhosSTOP Phosphatase Inhibitor Cocktail (Roche). Protein concentrations were determined using the BCA Protein Assay Kit (Thermo Scientific, #23227). Equal amounts of protein (10 µg per sample) were separated by SDS–PAGE on NuPAGE 4–12% BisTris gels (Thermo Scientific) and transferred onto nitrocellulose membranes. Membranes were blocked in TBS containing 0.05% Tween20 (TBST) and 5% skim milk for 1 h at room temperature, followed by incubation with primary antibodies diluted in TBST supplemented with 5% BSA overnight at 4 °C. After three 10min washes in TBST, membranes were incubated with horseradish peroxidase-conjugated secondary antibodies in blocking solution for 1 h at room temperature. Following three additional 10min washes in TBST, bands were visualized using chemiluminescent HRP substrate (Millipore, #WBKLS0100). Membranes were imaged on an iBright imaging system (Invitrogen, #FL1000) and quantified using ImageJ/Fiji (v1.53). Target protein signals were normalized to the total protein levels detected by Ponceau S staining on the corresponding membrane. Primary antibodies were used as follows: rabbit anti-p21^Waf1/Cip1^ (1:1000, Proteintech, #10355-1-AP), mouse anti-BCL2 (1:1000, Proteintech,#60178-1-Ig), rabbit anti-Cytochrome C (1:2000, Proteintech,#10993-1-AP), mouse anti-Caspase-3/p17/p19 (1:1000, Proteintech, #66470-2-Ig), rabbit anti-HSP60 (1:2500, Proteintech,#15282-1-AP), rabbit anti-COX4 (1:2500, Proteintech,#11242-1-AP), rabbit anti-VDAC1 (1:2500, Proteintech, #81538-1-RR), rabbit anti-TOMM20 (1:2000, Proteintech,#11802-1-AP). Secondary antibodies comprised mouse-anti rabbit horseradish peroxidase conjugates (Millipore, #AP188P, 1:2500 dilution) and goat anti-mouse horseradish peroxidase conjugates (Millipore, #AP181P, 1:2500 dilution).

### 5.10 Annexin V and Propidium Iodide Staining

Cell apoptosis was assessed using the 488Annexin V/PI Apoptosis Kit (Proteintech, #PF00005). Following Dox treatment, cells were washed twice with PBS and incubated in 1× Annexin V binding buffer containing 488Annexin V and PI for 15 min at room temperature in the dark. After staining, cells were washed twice with 1× Annexin V binding buffer and subsequently maintained in fresh 1× binding buffer for imaging. Fluorescence images were acquired using an EVOS M5000 inverted microscope (Thermo Fisher Scientific) equipped with 10× and 40× objectives.

### 5.11 Measurement of Mitochondrial bioenergetics

Cellular bioenergetics were assessed by measuring oxygen consumption rate (OCR) and extracellular acidification rate (ECAR) using the XFe96 Flux Analyzer (Agilent Seahorse XFe96 Analyzer). The Seahorse XF Cell Mito Stress Test Kit was used to evaluate mitochondrial function according to the manufacturer’s protocol. Cells were seeded at a density of 50,000 cells per well of XFe96 seahorse culture plates and incubated at 37 °C in a non-CO_2_ incubator for 1 h. At the end of incubation, cells were washed and culture media was switched to Seahorse Base Medium (XF DMEM medium, Agilent technologies, #103575) supplemented with 10 mM glucose, 2 mM L-glutamate, and 1 mM pyruvate.

Three repetitions of OCR/ECAR measurements were obtained and averaged for each assay stage (baseline, oligomycin, FCCP, Rotenone/antimycin A). Each measurement cycle comprised a 3-minute mixing phase followed by a 3-minute data acquisition period. Mitochondrial parameters including basal respiration, ATP-linked respiration, and spare respiratory capacity were determined by sequential injection of 1.5 µM oligomycin (ATP synthase inhibitor), 0.5 µM FCCP (carbonyl cyanide4[trifluoromethoxy]phenylhydrazone; an uncoupling agent), and a combination of 0.5 µM rotenone and antimycin A (complex I and complex III inhibitors, respectively). The mitochondrial respiration was analyzed using Agilent Seahorse Wave Pro software and Mito Stress Test Report Generator.

ATP-linked mitochondrial respiration was inferred from the oligomycin-induced reduction in OCR, whereas spare respiratory capacity was calculated as the difference between FCCP stimulated OCR and basal OCR. All OCR and ECAR values were normalized to protein content quantified using the BCA Protein Assay Kit (Thermo Scientific, #23227).

### 5.12 Statistics and data analysis

Statistical analyses were performed using GraphPad Prism v8.4.3 and v11. Data were presented as mean + SEM. Data normality was assessed using the Shapiro-Wilk and Kolmogorov Smirnov tests. Statistical differences between two groups were analyzed using an unpaired two-tailed Student’s *t*-test when data were normally distributed. Differences were defined as statistically significant at *p* < 0.05. Figures in the paper were prepared using Inkscape.

## Author Contributions

E.K., B.P., S.J. and N.V.J. conceived the study and E.K., N.V.J., M.Z.K.A, O.S., and B.P. designed the experiments. E.K., N.V.J., M.Z.K.A., and O.S. wrote the manuscript with input from all authors. N.V.J. performed 2D NPC culture experiments and associated downstream analyses. M.Z.K.A. conducted RNA-sequencing, bioinformatic analyses and data visualization. O.S. carried out hippocampal organoids culture and downstream analyses.

## Acknowledgements

We thank the Kang and Platt labs for their support and insightful discussions. We gratefully acknowledge the Microscopy and Histology Core Facility staff at the University of Aberdeen for their technical assistance.

## Data Availability Statement

All data are available in the main text or the supplementary information. RNA-sequencing data have been deposited in the European Nucleotide Archive (ENA) under accession number PRJEB111769.

