## supplementary information for "DNA damage-induced senescence reshapes transcriptomic and functional landscape of human neural progenitor cells"

**The PDF file includes:**

Supplementary Figures 1 to 5

Supporting data Figure 1

**Other Supplementary Information for this manuscript includes the following:**

Supplementary Tables 1 and 2

Source Data

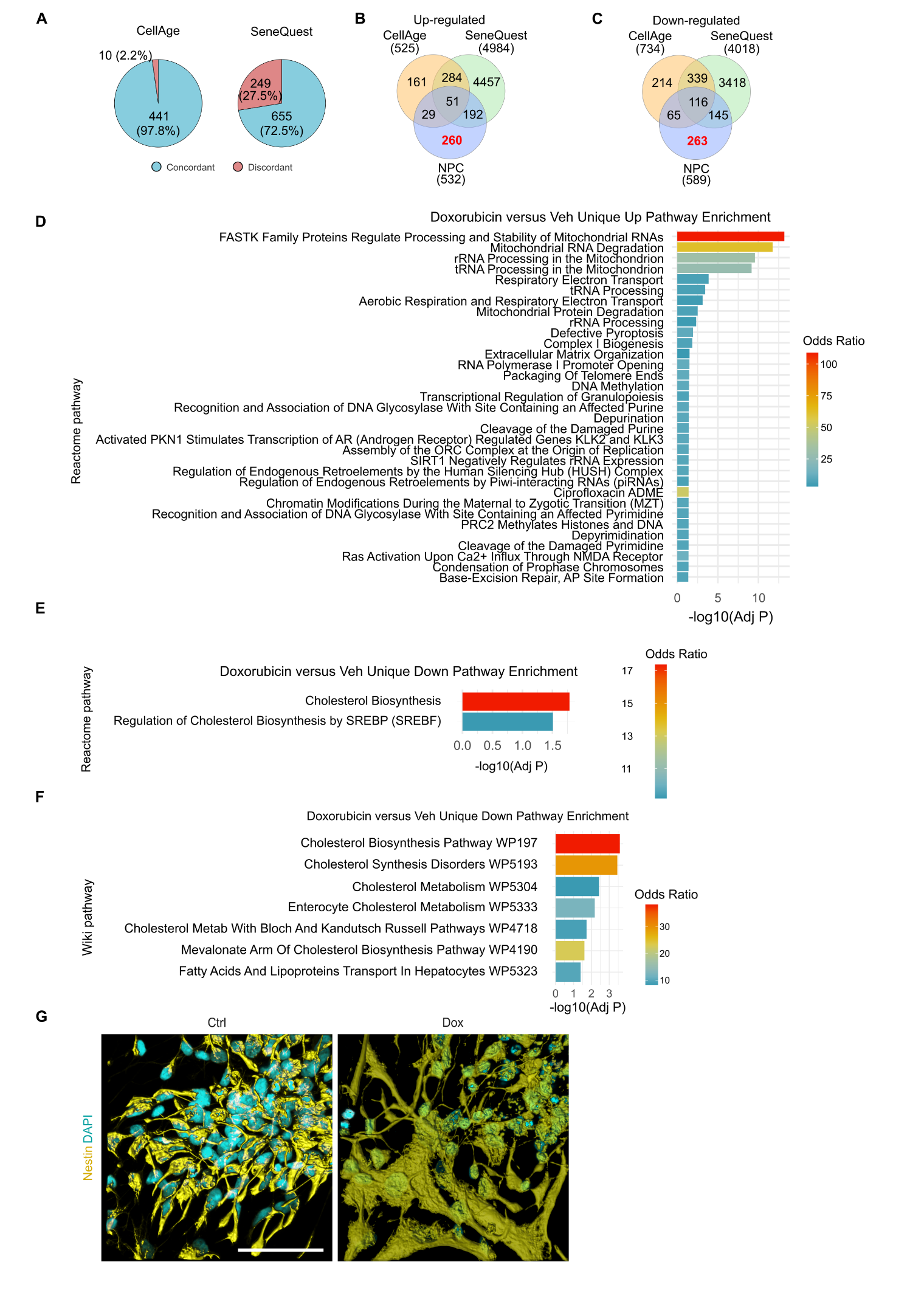

**Supplementary Fig. 1: Unique transcriptional signatures of Dox‑induced senescence in NPCs**

**(A)** Pie charts showing the number and percentage of genes with concordant or discordant expression changes between Dox‑treated NPC differentially expressed genes (DEGs) and senescence‑associated DEGs. Charts depict overlap with the CellAge database (left) and the SeneQuest database (right).

**(B)** Venn diagram comparing genes upregulated in Dox‑treated NPCs with the CellAge and SeneQuest senescence databases. Of 532 upregulated genes, 260 were unique to NPCs, not previously reported in senescence signature datasets.

**(C)** Venn diagram comparing genes downregulated in Dox‑treated NPCs with CellAge and SeneQuest. Of 589 downregulated genes, 263 were unique to NPCs, not present in senescence signature databases.

**(D)** Functional enrichment analysis of the 260 uniquely upregulated genes. Reactome pathways with FDR‑adjusted *p* < 0.05 are shown.

**(E)** Functional enrichment analysis of the 263 uniquely downregulated genes using Reactome pathways (FDR‑adjusted *p* < 0.05).

**(F)** Functional enrichment of the same 263 uniquely downregulated genes using WikiPathways (FDR‑adjusted *p* < 0.05).

**(G)** 3D reconstruction of z‑stack confocal images (Imaris) of control and Dox‑treated NPCs. Nestin immunostaining highlights senescence‑associated morphological changes, including increased cell‑body size, broader cell spreading, and irregular surface topology compared with the compact, uniform morphology of control NPCs. *Scale bar: 50 µm.*

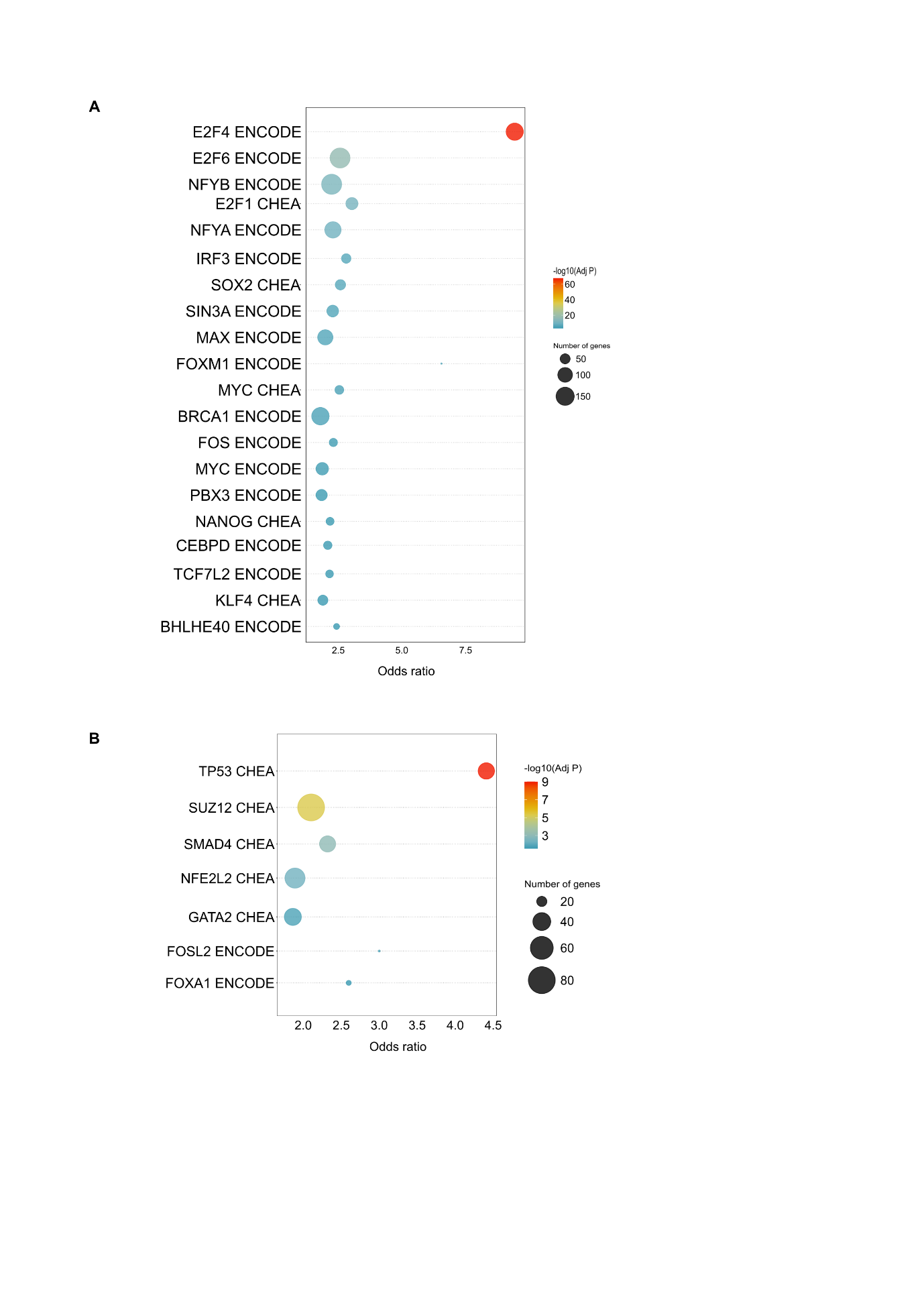

**Supplementary Fig. 2: Transcription factors regulating differentially expressed genes in Dox‑treated NPCs**

**(A–B)** Bubble plots showing transcription factors (TFs) significantly enriched among downregulated **(A)** and upregulated **(B)** genes in Dox‑treated NPCs. TFs with an FDR‑adjusted *p* < 0.05 identified using the consensus ENCODE and ChEA databases in Enrichr are displayed. The x‑axis shows the odds ratio (Enrichr), bubble size reflects the number of target genes regulated by each TF, and bubble colour represents the −log₁₀(FDR‑adjusted *p*).

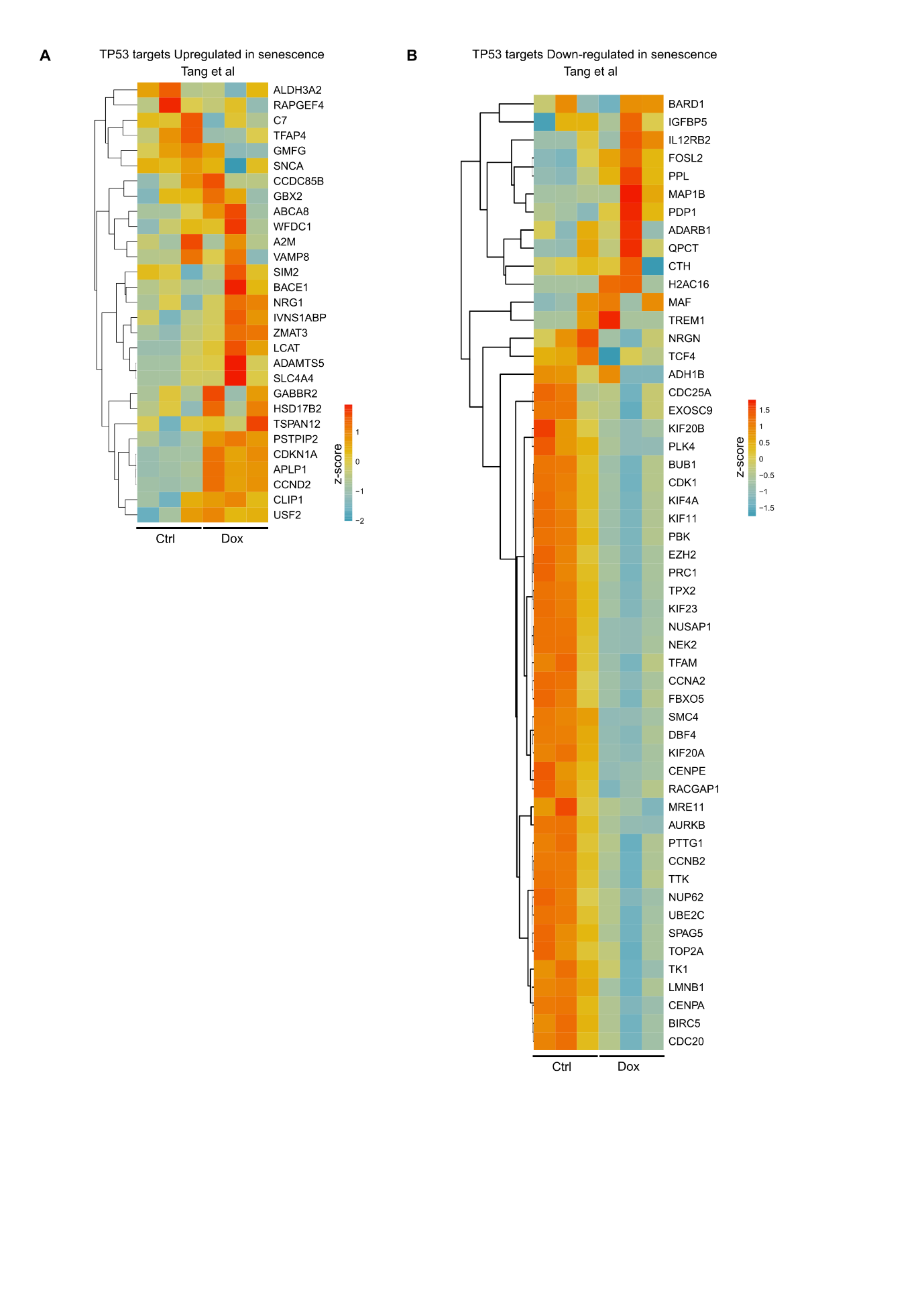

**Supplementary Fig. 3: Concordant regulation of previously reported TP53 target genes in Dox‑induced NPC senescence**

**(A–B)** Heatmaps showing expression patterns of TP53 target genes previously identified by *Tang et al.* as upregulated **(A)** or downregulated **(B)** during senescence. Each column represents an individual sample (three control and three Dox‑treated NPCs). Heatmaps were generated using VST‑transformed, normalized gene‑expression values, and colours represent the z‑score of expression for each gene across samples.

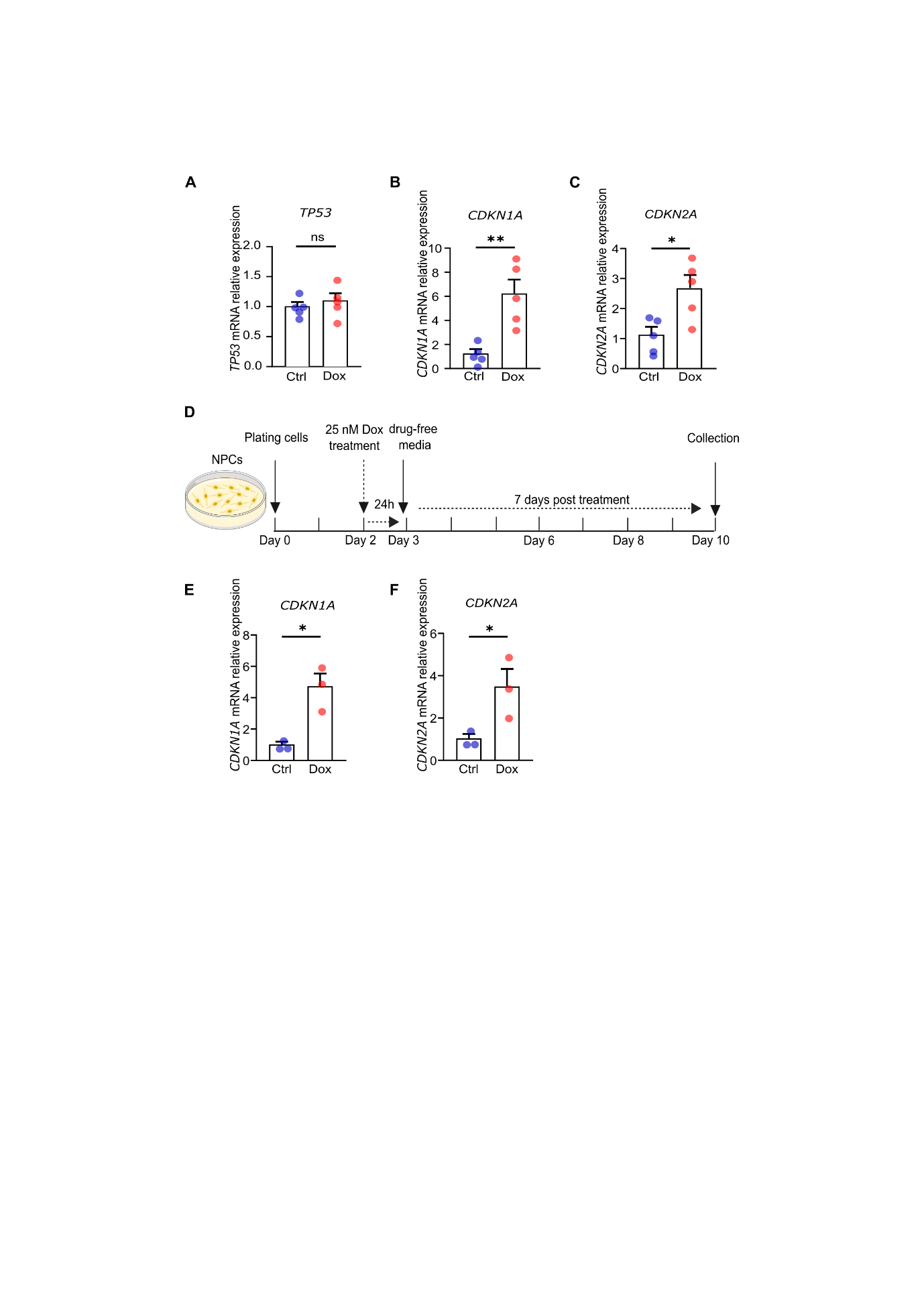

**Supplementary Fig. 4: qPCR validation of p53–p21–p16 pathway activation and persistence of senescence after Dox withdrawal**

(A–C) Relative mRNA expression levels of TP53 (p53) (A), CDKN1A (p21) (B), and CDKN2A (p16) (C) in Ctrl and Dox-treated‑NPCs measured by RT–qPCR. Values represent mean+ SEM from n = 5 biological replicates, normalized to the housekeeping genes RPS18 and 18S, **p < 0.05*; ***p < 0.01*; ns: not significant, unpaired two-tailed Student's t test.

**(D)** Schematic of the experimental paradigm used to assess senescence stability after drug withdrawal. NPCs were treated with 25 nM Dox for 24 h; Dox was removed on day 3, and cells were maintained in drugfree medium for 7 additional‑ days. Cells were harvested on day 10 for transcriptional analysis.

**(E–F)** Relative mRNA expression levels of *CDKN1A* (p21) (E) and *CDKN2A* (p16) (F) following the 7-day drug‑-free interval, measured by RT–qPCR‑. Values represent mean± SEM from n = 3 biological replicates, normalized to the housekeeping genes RPS18 and 18S, **p < 0.05*; ns: not significant, unpaired two-tailed Student's t test.

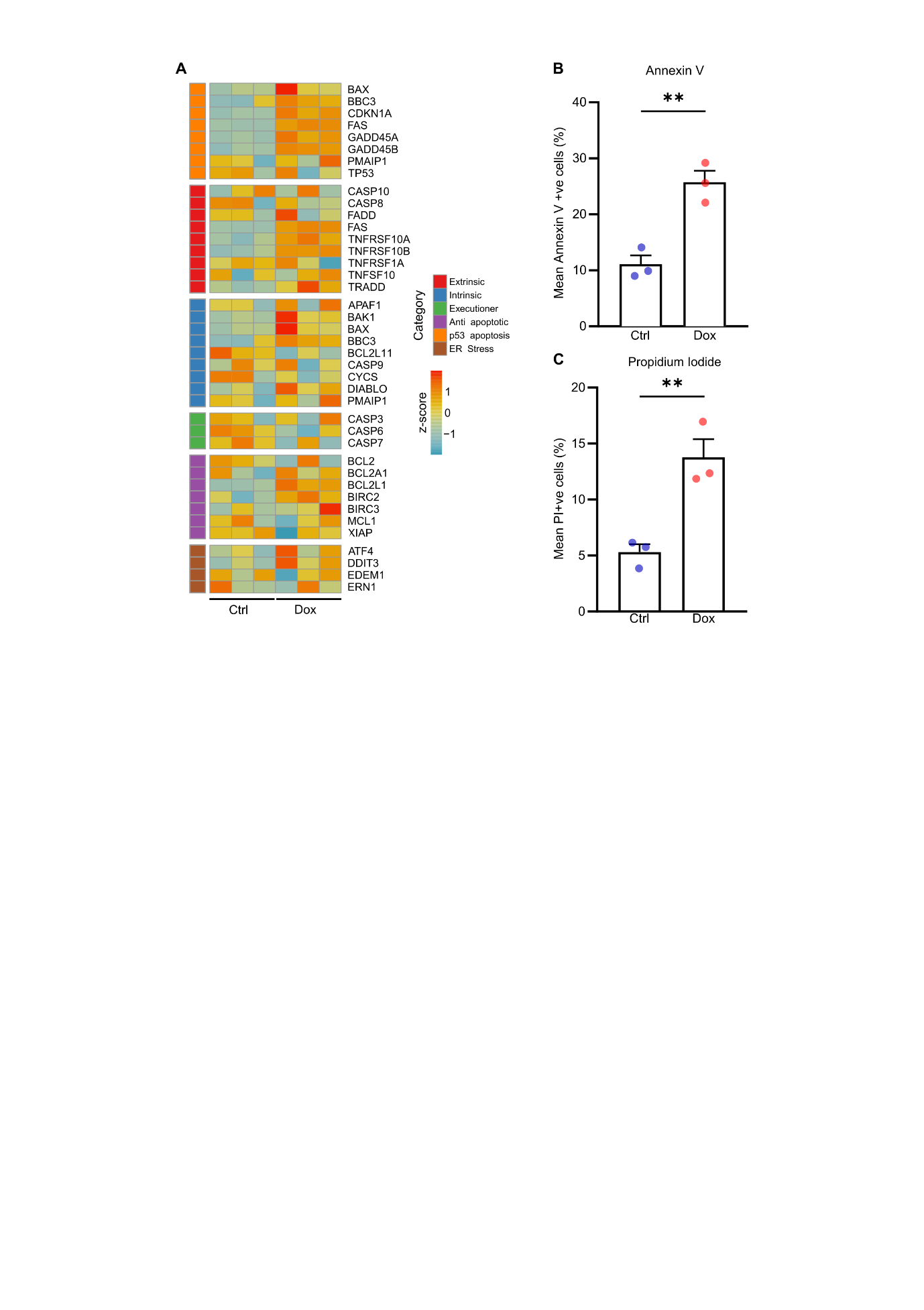

**Supplementary Fig. 5: Senescent NPCs exhibit apoptosis profiles comparable to controls**

**(A)** Heatmaps showing the expression patterns of apoptosis‑related genes, grouped by pathway category. Each column represents an individual sample (three control and three Dox‑treated NPCs). Heatmaps were generated from VST‑transformed, normalized gene‑expression values, and colours indicate the z‑score of each gene across samples.

**(B)** Quantification of Annexin V+(B) cells in Ctrl and Dox-treated NPCs, normalized to total cell count. Values represent mean ± SEM from *n* = 3 biological replicates, ***p < 0.01*, unpaired two-tailed Student's t test.

**(C)** Quantification of PI+ cells in Ctrl and Dox-treated NPCs, normalized to total cell count. Values represent mean ± SEM from *n* = 3 biological replicates, ***p < 0.01*, unpaired two-tailed Student's t test.

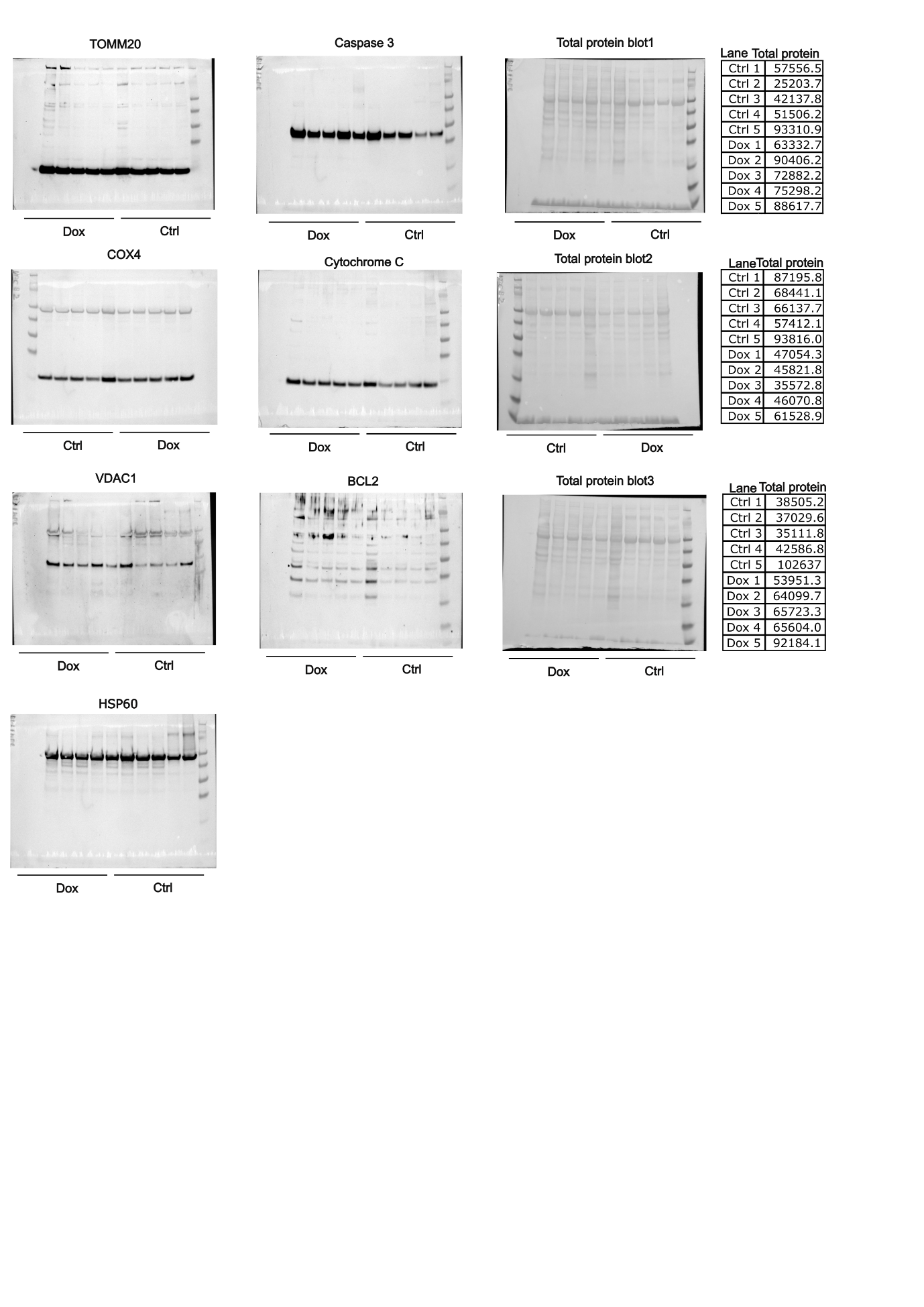

**Supporting data Fig.1. Full-length uncropped Western blot images corresponding to main figures**
